# A patient-derived amyotrophic lateral sclerosis blood-brain barrier cell model reveals focused ultrasound-mediated anti-TDP-43 antibody delivery

**DOI:** 10.1101/2024.02.22.581567

**Authors:** Joanna M. Wasielewska, Mauricio Castro Cabral-da-Silva, Martina Pecoraro, Tam Hong Nguyen, Vincenzo La Bella, Lotta E. Oikari, Lezanne Ooi, Anthony R. White

## Abstract

**Background:** Amyotrophic lateral sclerosis (ALS) is a rapidly progressing neurodegenerative disorder with minimally effective treatment options. An important hurdle in ALS drug development is the non-invasive therapeutic access to the motor cortex currently limited by the presence of the blood-brain barrier (BBB). Focused ultrasound and microbubble (FUS^+MB^) treatment is an emerging technology that was successfully used in ALS patients to temporarily open the cortical BBB. However, FUS^+MB^-mediated drug delivery across ALS patients’ BBB has not yet been reported. Similarly, the effects of FUS^+MB^ on human ALS BBB cells remain unexplored.

**Methods:** Here we established the first FUS^+MB^-compatible, fully-human ALS patient-cell-derived BBB model based on induced brain endothelial-like cells (iBECs) to study anti-TDP-43 antibody delivery and FUS^+MB^ bioeffects *in vitro*.

**Results:** Generated ALS iBECs recapitulated disease-specific hallmarks of BBB pathology, including changes to BBB integrity, permeability and TDP-43 proteinopathy. Our results also identified differences between sporadic ALS and familial (*C9orf72* expansion carrying) ALS iBECs reflecting patient heterogeneity associated with disease subgroups. Studies in these models revealed successful ALS iBEC monolayer opening *in vitro* with a lack of adverse cellular effects of FUS^+MB^. This was accompanied by the molecular bioeffects of FUS^+MB^ in ALS iBECs including changes in expression of tight and adherens junction markers, and drug transporter and inflammatory mediators, with sporadic and C9orf72 ALS iBECs generating transient specific responses. Additionally, we demonstrated an effective increase in the delivery of anti-TDP-43 antibody with FUS^+MB^ in C9orf72 (2.7-fold) and sporadic (1.9-fold) ALS iBECs providing the first proof-of-concept evidence that FUS^+MB^ can be used to enhance the permeability of large molecule therapeutics across the BBB in a human ALS *in vitro* model.

**Conclusions:** Together, our study describes the first characterisation of cellular and molecular responses of ALS iBECs to FUS^+MB^ and provides a fully-human platform for FUS^+MB^-mediated drug delivery screening on an ALS BBB *in vitro* model.

## BACKGROUND

Amyotrophic lateral sclerosis (ALS) is a neurodegenerative disorder characterised by the gradual loss of motor neurons (1). Despite major progress in ALS drug discovery, the effective translation of preclinical therapeutic agents to their clinical application in ALS patients remains limited, with approved therapeutics offering only a modest increase in life expectancy (1,2).

ALS pathophysiology is complex, with multiple pathways converging into upper and lower motor neuron degeneration (1). Correspondingly, extensive evidence suggests early pathological changes in the upper motor neurons in the motor cortex supporting the development of motor cortex-directed therapies (3–9). However, targeting cortical neurons provides challenges due to the presence of the blood-brain barrier (BBB), which markedly limits the penetration of ALS therapeutics into the central nervous system (CNS) (2,10–14).

Additionally, evidence from animal and human studies indicates BBB dysfunction in ALS, and suggests cerebrovascular involvement in disease development and progression (15–18). Changes in BBB ultrastructure and barrier integrity were found in preclinical models and in patients, leading to increased BBB permeability and infiltration of blood-derived proteins, erythrocytes and immune cells into the brain (16,17,19–24). Others reported degeneration of BBB-forming cells including brain endothelial cells (BECs), pericytes and astrocytes, and found impaired expression of tight and adherens junctions in BECs (19,25,26). In addition, alterations in BBB drug transporters such as P-glycoprotein (P-pg) and breast cancer resistance protein (BCRP) were identified in ALS (13,27). Collectively, these changes further contribute to the complex interaction of drugs with pathologically altered BBB limiting their clinical effectiveness. Thus, non-invasive therapeutic access to the motor cortex still poses a critical challenge in ALS drug development.

Focused ultrasound combined with activated microbubbles (FUS^+MB^) is an innovative method facilitating temporary BBB opening, therefore enabling targeted CNS drug delivery (28). In animal studies, the application of FUS^+MB^ aided in the delivery of ALS therapeutics across the BBB (29–31), providing additional therapeutic benefit as compared to the drug alone (30,31). In the first-in-human clinical study, FUS^+MB^ induced a transient increase in the leakage of contrast agent across the BBB in the motor cortex of four patients demonstrating the safety and feasibility of BBB opening in ALS (32). However, FUS^+MB^-mediated ALS drug delivery has not yet been achieved in humans. Similarly, the molecular effects of FUS^+MB^ on human ALS BBB cells are currently unknown.

Here we developed a novel ALS patient-cell-derived *in vitro* platform to study the bioeffects of FUS^+MB^ on human BECs and investigated FUS^+MB^-enhanced antibody transport across ALS BBB *in vitro.* To accurately model the heterogeneity of the ALS BBB pathology (16) we utilised human induced pluripotent stem cells (iPSCs) from a sporadic ALS patient and those from an individual carrying the intronic *C9orf72* hexanucleotide expansion (> 30 GGGGCC) that is found in an estimated 40% of familial ALS cases (33), and differentiated them towards brain endothelial-like cells (iBECs). Our results demonstrated phenotypical differences in ALS iPSC-derived iBECs as compared to healthy donor controls, as well as between sporadic and familial ALS cells, replicating some of the key features of ALS BBB pathology found in humans (24,26). We also presented a lack of adverse effects of FUS^+MB^ on ALS iBECs viability and the reversibility of barrier opening *in vitro*, lending further support to the clinical safety of FUS^+MB^ in ALS patients. Finally, we achieved improved delivery of anti-TDP-43 antibody with FUS^+MB^ demonstrating the applicability of FUS^+MB^ to enhance large molecule therapeutic permeability in human ALS BBB *in vitro*.

Together this proof-of-concept study provides the first ALS patient-cell-derived BBB model for FUS^+MB^-enhanced therapeutic permeability screening, thereby contributing to the translational advancement of ultrasound-based therapies in ALS.

## MATERIALS AND METHODS

### Human iPSCs generation and characterisation

Three human iPSC lines were utilised in this study (**Table S1**). The healthy donor (HD) control line HDFa was previously generated by prof. Jose M. Polo laboratory (Monash University) and characterised by us (34,35). Two ALS iPSC lines were generated from patients’ skin fibroblasts. A punch biopsy was obtained from the left arm of two patients clinically diagnosed with either sporadic or familial (*C9orf72*; > 30 GGGGCC repeats) ALS (**Table S2**) after informed consent. The obtained biopsy was immediately placed in cold phosphate-buffered saline (PBS). The skin explant was then cut into pieces, which were cultured in Dulbecco’s modified Eagle’s medium supplemented with 10% calf serum, 2 mM L-glutamine, 5 mM pyruvate, 100 U/mL penicillin and 100 μg/mL streptomycin. Fibroblasts were maintained in culture and then frozen in liquid nitrogen. The respective ALS iPSC lines IT_002-04 and IT_004-04 were generated from the patient’s fibroblasts using repeated mRNA transfection for pluripotency transcription factors OCT4, KLF4, SOX2, c-MYC, LIN28 and NANOG as described previously (36). The STR profiling was performed and confirmed 100% match to the donor fibroblasts.

iPSCs from all lines were cultured in StemFlex^TM^ medium on human recombinant vitronectin (Thermo Fisher Scientific) as described (34,35,37). All iPSC lines were characterised for nuclear expression of stemness markers Nanog and SOX2 by immunofluorescence as we previously described (34,35).

### Differentiation and characterisation of human induced brain endothelial-like cells (iBECs)

Induced brain endothelial-like cells (iBECs) were generated from human iPSCs as previously described (34,35,37,38). Briefly, iPSCs were plated at 20,000 cells/cm^2^ on human embryonic stem cell (hESC)-qualified Matrigel (Corning) coating in StemFlex^TM^ medium supplemented with 10 μM Rho-associated kinase inhibitor (iROCK) to initiate differentiation (34,35,37). Cells were then cultured for six days in an unconditioned medium consisting of DMEM/F12+GlutaMAX (Life Technologies), 20% KnockOUT serum replacement (Life Technologies), 1 x non-essential amino acids (Life Technologies) and 0.1 mM β-mercaptoethanol (Sigma) (34,37,38), and subsequently for 2 days in endothelial cell medium (EC; Life Technologies) supplemented with 2% B27 (Life Technologies), 20 ng/ml basic fibroblast growth factor (FGFb; Peprotech) and 10 μM retinoic acid (RA) (34,37,38). Next, cells were singularised with StemPro^TM^ Accutase (Gibco) and purified on collagen IV from human placenta (Sigma) and human plasma fibronectin (Life Technologies) coated plastic culture plates or 0.4 µm or 3.0 µm pore Transwell inserts (Corning) as described previously (34,35,37,38). After one day, the medium was changed to EC+B27 (without FGFb and RA) to allow full cell maturation. All assays were performed 48 h following cell purification and iBECs were cultured in EC+B27 under normoxia (normoxia conditions (37 °C, 5% CO2) for the duration of experimental assays (34,37,38).

Differentiated iBECs were characterised for expression of BEC-specific markers occludin, claudin-5, ZO-1 and Glut-1 by immunofluorescence as we previously described (34,35,37) utilising antibodies listed in **Table S3**.

To assess iBECs barrier integrity, cells were cultured on 24-well Transwell insert with 0.4 µm pore or 3.0 µm pore membrane (Corning) and transendothelial electrical resistance (TEER) across iBEC monolayer was measured with the EVOM3 Volt/Ohmmeter (World Precision Instruments) as we previously described (34,37). TEER was measured in three areas per Transwell and the resistance of the blank (no-cells) Transwell was subtracted before averaging.

To assess the passive permeability of iBEC monolayer, cells were cultured in 0.4 µm or 3.0 µm pored Transwell inserts and fluorescein isothiocyanate (FITC)-conjugated dextran molecules of 3– 5 kDa or 150 kDa (Sigma) were added at 0.5 mg/ml to the top chamber of the Transwell insert for 24 h. Next, fluorescence signal intensity (490 nm excitation/520 nm emission) was measured in cell culture medium collected from the top and bottom chamber of the Transwell system (three technical replicates per Transwell) using a fluorescent plate reader (Biotek Synergy H4) and clearance volume calculated as previously described (37,38).

### TDP-43 protein expression analysis

For immunofluorescence-based quantification of TDP-43 expression iBECs were cultured on collagen IV and fibronectin-coated plastic coverslips and fixed with 4% paraformaldehyde (PFA) for 15 min at room temperature (RT). Next, cells were washed with PBS and permeabilised with 0.3% Triton X-100 for 10 min and blocked for 1 h at RT with 2% bovine serum albumin (BSA, Sigma)/2% normal goat serum (GS, Chemicon) in PBS. Primary antibodies for TDP-43 (1:200) and ZO-1 (1:100) (**Table S3**) were diluted in a blocking solution and incubated with cells overnight at 4°C. Then cells were washed with PBS and incubated with secondary antibodies (**Table S3**) diluted at 1:250 in a blocking solution for 1 h at RT. Subsequently, cells were washed with PBS and nuclear counterstaining was performed with Hoechst (1:5000). Coverslips were mounted with ProLong Gold Antifade (Invitrogen) and imaged at 100x magnification and identical image acquisition setting by the investigator blinded to cell genotype using a Zeiss 780 laser scanning confocal microscope. Antibody specificity was confirmed by performing secondary antibody-only controls.

Cytoplasmic and nuclear TDP-43 signal intensity (AU) in iBECs was measured in acquired images with ImageJ software, using methodology adapted from Quek *et al.* 2022 (39) where ZO-1 and Hoechst-stained cell images were used to create a mask for whole-cell and nucleus, respectively. Image contrast and brightness were uniformly increased using ZEN Black Software (Zeiss) for presentation purposes.

### FUS^+MB^-mediated delivery of 150 kDa dextran and anti-TDP-43 antibody

For ultrasound-mediated permeability studies, iBECs were cultured in Transwell inserts with 3.0 µm pores and exposed to FITC-conjugated 150 kDa dextran (0.5 mg/ml) or anti-TDP-43 antibody (1 µM, Sigma, Cat#T1705) and 10 µl of phospholipid-shelled microbubbles with octafluoropropane gas core prepared in-house as described in (40). Wells were then immediately exposed to FUS at 286 kHz center frequency, 0.3 MPa peak rarefactional pressure, 50 cycles/burst, burst period 20 ms, and a 120 s sonication time as we previously described (34,37,41). 24 h following the treatment, cell culture medium was collected for dextran fluorescence signal intensity assessment as described above. Anti-TDP-43 antibody concentration was determined by Rabbit IgG enzyme-linked immunoassay kit (ELISA, Abcam) following manufacturer’s instructions. For FUS^+MB^ studies, fold change in detected dextran clearance volume or antibody concentration was calculated relative to its untreated (UT) control for each line at 24 h.

### Monolayer morphology assessment post FUS^+MB^

To investigate potential mechanical damage of iBEC monolayer following FUS^+MB^ cells were cultured on collagen IV and fibronectin-coated plastic coverslips and exposed to FUS at the parameters described above and 20 µl MB. Immediately and 24 h post-treatment cells were fixed with 4% PFA for 15 min and immunofluorescence staining for ZO-1 was performed as described above. Coverslips were mounted with Dako Mounting medium (Agilent) and imaged at 20x magnification by the investigator blinded to the treatment condition using a Zeiss 780 confocal microscope.

### Lactate dehydrogenase (LDH) viability assay

The effects of FUS^+MB^ on iBECs viability were assessed utilising CyQUANT LDH Cytotoxicity Assay (Thermo Fisher Scientific) as we previously described (37). Briefly, cells were cultured in 24-well plates and exposed to FUS at the parameters described above and 20 µl MB and cell culture media samples were collected immediately (30 min) and 24 h post-treatment. The levels of LDH enzyme were assessed with CyQUANT assay kit following manufacturer’s instructions and calculated relative to its untreated (UT) control for each line (37).

### iBEC monolayer integrity assessment post FUS^+MB^

To assess the effects of FUS^+MB^ on iBEC monolayer integrity, iBECs were cultured on 3.0 µm pore Transwells and exposed FUS at the parameters described above and 10 µl MB. TEER was measured immediately (imm.; 1 h) and 24 h post-treatment in three areas per Transwell as described above and in (37). Fold change in TEER compared to respective untreated (UT) controls was calculated.

### RNA extraction, cDNA synthesis and quantitative real-time PCR (qPCR)

For cell phenotype characterisation, iPSC and iBEC were cultured under normal conditions and remained untreated prior to RNA collection. For FUS^+MB^ effects studies, cells were cultured in 24-well plates and exposed to FUS at the parameters described above and 20 µl MB 30 min (immediately; imm.) and 24 h prior to RNA collection as we described (37).

For RNA collection, cells were rinsed with PBS and lysed in TRIzol^TM^ reagent (ThermoFisher Scientific) and total RNA was extracted using the Direct-zol RNA Miniprep Kit (Zymo Research) according to the manufacturer’s instructions. Isolated RNA quality and quantity was measured using NanoDrop^TM^ Spectrophotometer. For quantitative real-time polymerase chain reaction (qPCR) studies, 150 ng of total RNA was converted to complementary DNA (cDNA) using SensiFAST^TM^ cDNA synthesis kit following manufacturer instructions (Bioline) and qPCR performed in technical triplicate using SensiFAST™ SYBR® Lo-ROX Kit following manufacturer instructions (Bioline) on QuantStudio^TM^ 5 Real-Time PCR system as previously described (34,35,37,41). Obtained Ct values were normalised by the Ct values of *18S* endogenous control (ΔCt). Housekeeping gene expression of *18S* was found to be consistent across cell lines. The ΔΔCt values were calculated as 2(^-ΔCt^) and multiplied by 10^6^. Technical triplicates were averaged per sample and log transformed for statistical analysis. Utilised primer sequences are presented in **Table S4**.

### Statistical analysis

Statistical analysis was performed using GraphPad Prism version 9.4.0. For a two-group comparison with normal distribution, *F* test of equality of variances was performed and data was analysed with unpaired Student’s *t*-test (two-tailed; data with equal variances) or unpaired Welch’s *t*-test (two-tailed; data with unequal variances). When comparisons between three or more groups were analysed, one-way ANOVA followed by post-hoc tests were used. *P* < 0.05 was considered statistically significant. Z-scores were calculated and values with Z-scores above or below two standard deviations (SD) of the mean were identified as outliers and excluded from analysis. Results are shown as mean ± SEM. The number of independent (*n*) replicates used for each experiment is specified in figure legends. The number of technical replicates included in each assay is stated in the respective materials and methods sections.

## RESULTS

### ALS iBECs demonstrate an altered phenotype compared to healthy donor controls

To study the ALS BBB *in vitro*, we generated human iBECs from a healthy donor (HD) iPSC line previously characterised by us (34,35) and two novel ALS iPSC lines derived from sporadic and familial (C9orf72 variant) ALS patients (**Table S1, S2**). The stemness of utilised iPSC lines was ascertained by the nuclear expression of SOX2 and Nanog proteins and iPSC-like morphology was confirmed by the formation of characteristic uniform colonies with defined edges (**Figure S1A**). Expression of pluripotency-regulator genes *SOX2*, *NANOG* and *OCT4* (42) was also confirmed at mRNA level and demonstrated no differences between control and ALS iPSCs, suggesting phenotypic similarity of selected iPSC lines (**Figure S1B**). The iPSCs were differentiated towards iBECs following previously published protocols (34,35,37) and successful generation of cells with brain endothelial-like phenotype confirmed by the formation of cobblestone-like monolayers and expression of BEC-associated markers including occludin, claudin-5, zonula occludens-1 (ZO-1) and glucose transporter-1 (Glut1) (**Figure 1A**). When compared to parental iPSCs, mature iBECs overall presented decreased expression of pluripotency marker genes *SOX2*, *NANOG* and *OCT4* and increased expression of BEC-associated tight and adherens junction genes: occludin (*OCLN)*, claudin-5 (*CLDN5)*, ZO-1 (*TJP1)* and ve-cadherin (*CDH5)* (**Figure S2**), suggesting a switch towards the desired endothelial cell phenotype.

**Figure 1.**
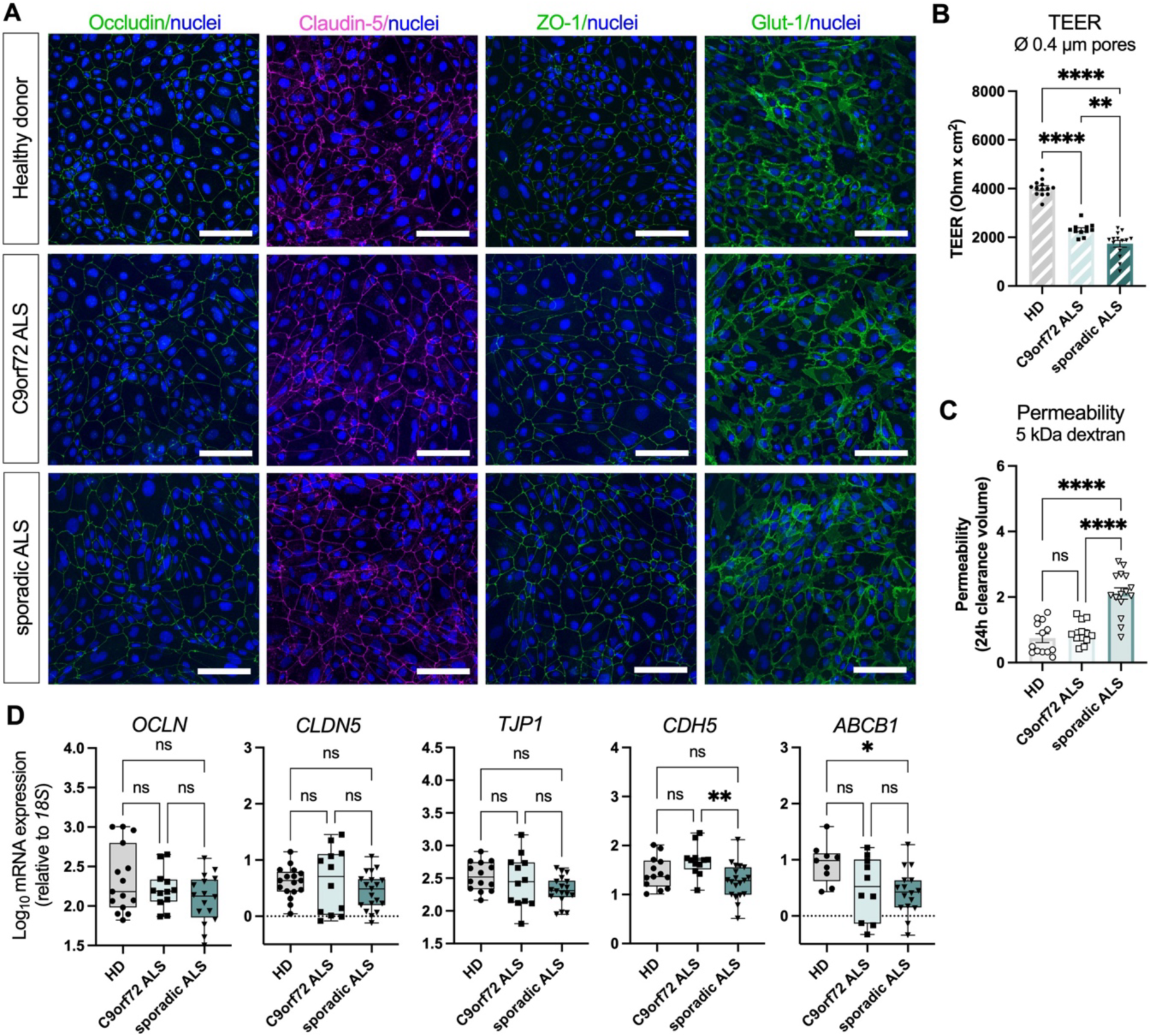
Characterisation of brain endothelial-like cells (iBECs) generated from control and ALS human induced pluripotent stem cells (iPSCs). (**A**) Representative immunofluorescence images of occludin (green), claudin-5 (magenta), ZO-1 (green) and Glut-1 (green) with Hoechst nuclear counterstaining (blue) in healthy donor, C9orf72 ALS, and sporadic ALS iBECs. Scale bar = 100 µm. (**B**) Trans-endothelial electrical resistance (TEER, Ohm x cm^2^) of a healthy donor, C9orf72 ALS, and sporadic ALS iBECs, measured in 0.4 µm pore Transwells. A minimum of *n* = 11 independent replicates per line. (**C**) Passive permeability of FITC-conjugated 5 kDa dextran in healthy donor, C9orf72 ALS, and sporadic ALS iBECs assessed in 0.4 µm pore Transwells. A minimum of *n* = 12 independent replicates per line. Data presented as clearance volume of dextran at 24 h. (**D**) Relative mRNA expression of brain endothelial cell marker genes *OCLN*, *CLDN5, TJP1*, *CDH5* and *ABCB1* transporter in healthy donor, C9orf72 ALS, and sporadic ALS iBECs. Data presented as Log10 of ΔΔCT x 10^6^, relative to *18S*. A minimum of *n* = 8 independent replicates per line. Data in (B-D) analysed with One-way ANOVA with Tukey’s test, error bars = SEM. \**P*<0.05, \*\**P*<0.01, \*\*\*\**P*<0.0001.

Since BBB pathology was previously described in ALS patients (15,16,24,26), we next examined the development of a disease-related phenotype in generated patient-cell derived iBECs. When cultured on traditionally used 0.4 µm pore Transwell inserts (34,38,43,44), cells from all lines formed a confluent monolayer with physiologically relevant (45,46) integrity measured by the transendothelial electrical resistance (TEER; healthy donor: 4024 ± 95.02, C9orf72 ALS: 2312 ± 79.3, sporadic ALS: 1733 ± 137.4 Ohm x cm^2^, mean ± SEM, **Figure 1B**). Similar to BBB integrity impairment observed in ALS *in vivo* (19,26), ALS iBECs formed a monolayer of reduced (*P*<0.0001) TEER compared to control cells, with sporadic ALS cells demonstrating lower (*P*<0.01) barrier integrity than C9orf72 expansion-carrying ALS iBECs (**Figure 1B**). We then assessed the passive permeability of iBEC monolayers to biologically inert 5 kDa dextran fluorescent tracer (37) and found its increased (*P*<0.0001) leakage in sporadic ALS iBECs compared to healthy donor and C9orf72 ALS cells (**Figure 1C**). This was accompanied by downregulation (*P*<0.01) of *CDH5* mRNA expression in sporadic ALS iBECs suggesting there may be changes to adherens junction complexes in these cells (**Figure 1D**); while the expression of tight junction markers *OCLN*, *CLDN5* and *TJP1* did not differ between control and ALS cells. Interestingly, the expression of *ABCB1* encoding for drug transporter P-gp was decreased (*P*<0.05) in sporadic ALS iBECs (**Figure 1D**), consistent with previous observations of P-gp dysregulation in ALS (13,27,47–49).

### ALS iBECs reveal cytoplasmic accumulation of TDP-43 protein

TAR DNA-binding protein 43 (TDP-43) proteinopathy is a hallmark of ALS (50,51) with TDP-43 protein aggregates recently being identified in cortical small blood vessels of sporadic ALS patients (52). To investigate the potential TDP-43-related vasculopathy (52) in our model, we evaluated the cellular localisation of TDP-43 protein in control and ALS iBECs by immunofluorescence. Our analysis revealed cytoplasmic accumulation of TDP-43 protein aggregates in ALS iBECs (Figure 2A, **Figure S3A**) with both C9orf72 (*P*<0.05) and sporadic (*P*<0.0001) ALS cells demonstrating higher cytoplasmic TDP-43 protein signal intensity compared to healthy donor controls (Figure 2B). Cytosolic TDP-43 aggregates were primarily granular in shape and more profound (*P*<0.05) in sporadic ALS as compared to C9orf72 ALS iBECs (Figure 2A-B, **Figure S3A**). Interestingly, healthy donor iBECs presented low (*P*<0.0001) TDP-43 protein expression in the nucleus (compared to ALS cells), with no difference found in nuclear TDP-43 expression between sporadic and C9orf72 iBECs (Figure 2B). Hence, to assess the distribution of total TDP-43 protein in studied cells, we examined the ratio of cytoplasmic to nuclear TDP-43 signal and identified its increase in sporadic ALS iBECs compared to healthy donor controls (*P*<0.05) and C9orf72 ALS iBECs (*P*<0.01, Figure 2C) corroborating TDP-43 mislocalisation in sporadic ALS cells. No signal was found in the secondary-only controls (**Figure S3B**) confirming the specificity of the utilised anti-TDP-43 antibody.

**Figure 2.**
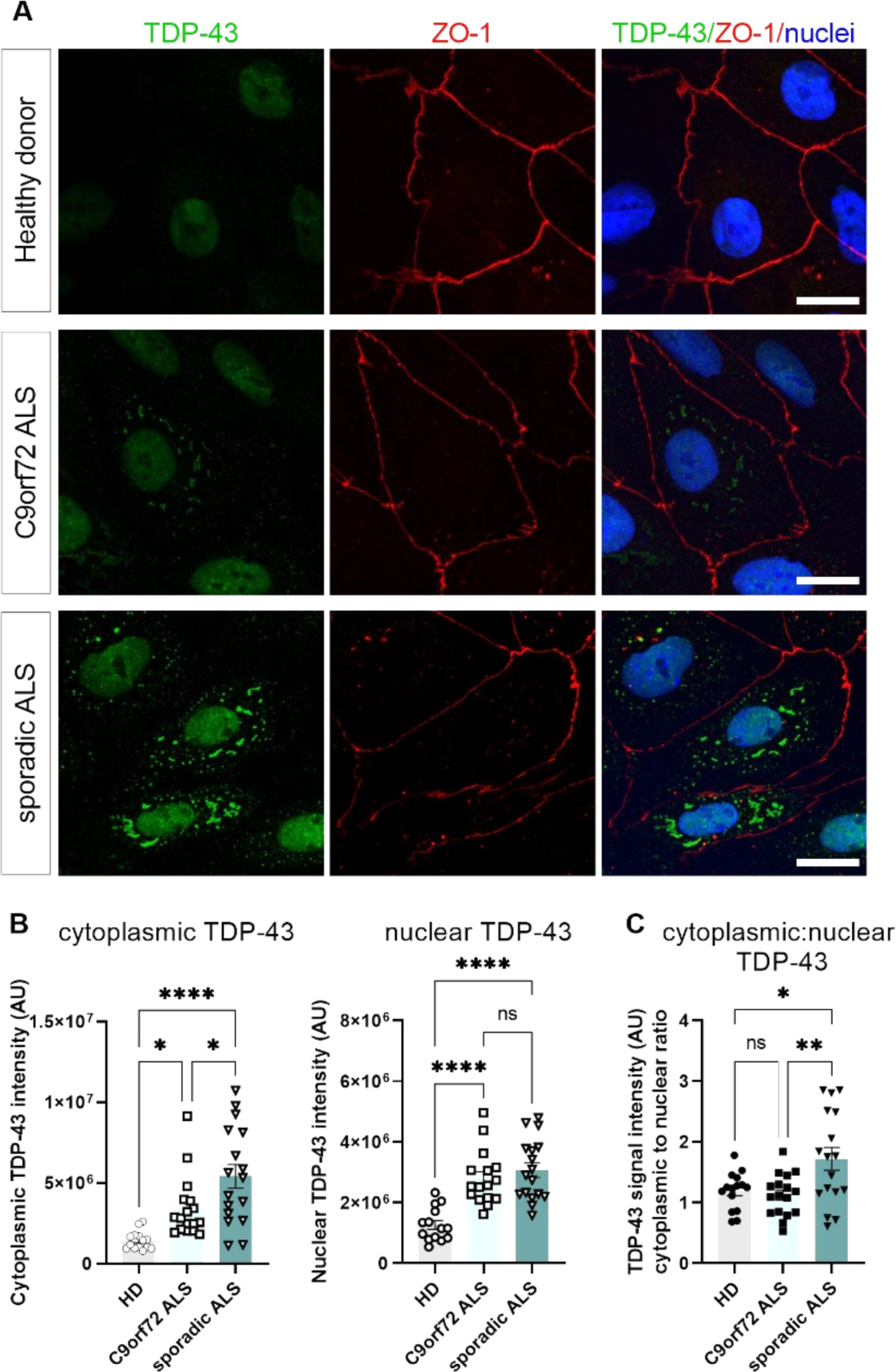
Expression of TDP-43 protein in control and ALS iBECs. Representative immunofluorescence images of TDP-43 (green) and ZO-1 (red) with Hoechst nuclear counterstaining (blue) in healthy donor, C9orf72 ALS, and sporadic ALS iBECs. Scale bar = 20 µm. (**B**) Comparison of TDP-43 signal intensity in the cytoplasm and nuclei of healthy donor, C9orf72 ALS, and sporadic ALS iBECs. (**C**) Comparison of cytoplasmic to nuclear TDP-43 signal intensity ratio in healthy donor, C9orf72 ALS, and sporadic ALS iBECs. A total of *n* = 14-17 cells analysed from four independent replicates per line in (B,C). Data in (B,C) analysed with One-way ANOVA with Tukey’s test \**P*<0.05, \*\**P*<0.01, \*\*\*\**P*<0.0001, error bars = SEM.

### FUS^+MB^ facilitates controlled iBEC monolayer opening in vitro

We have previously demonstrated that FUS^+MB^ applied at clinically relevant (32,53) parameters leads to reversible iBEC monolayer opening in familial and sporadic Alzheimer’s disease (AD) BBB models (34,37,41). In this study, we aimed to utilise characterised ALS patient-derived iBECs as a platform to investigate the FUS^+MB^-mediated delivery of large molecule therapeutics at the ALS BBB.

Our prior investigation revealed that most commonly used Transwell inserts with pores of 0.4 µm in diameter (34,38,43,44,54,55) are not suitable to reliably assess the permeability of large (≥150 kDa) molecules and instead, Transwells with membranes containing 3.0 µm pores provide a technically advantageous alternative (37). Hence, we first optimised ALS iBECs culture conditions leading to the development of a confluent cell barrier in Transwell inserts with 3.0 µm pores (Figure 3A). As expected (56), control and ALS iBECs generated a barrier with lower TEER when cultured on 3.0 µm pored membranes as compared to the 0.4 µm Transwell (healthy donor: 1121 ± 44.35, C9orf72 ALS: 845 ± 42.61, sporadic ALS: 673 ± 26.54 Ohm x cm^2^, mean ± SEM, Figure 3B), while reaching 500 Ohm x cm^2^ threshold established as a benchmark of effective *in vitro* barrier formation (57,58). Experiments performed in this Transwell format also replicated the reduced (*P*<0.0001) TEER in ALS iBECs compared to healthy donor cells, as well as a further decrease (*P*<0.01) in sporadic ALS iBECs TEER when compared to C9orf72 cells (Figure 3B), corroborating our findings (Figure 1B).

**Figure 3.**
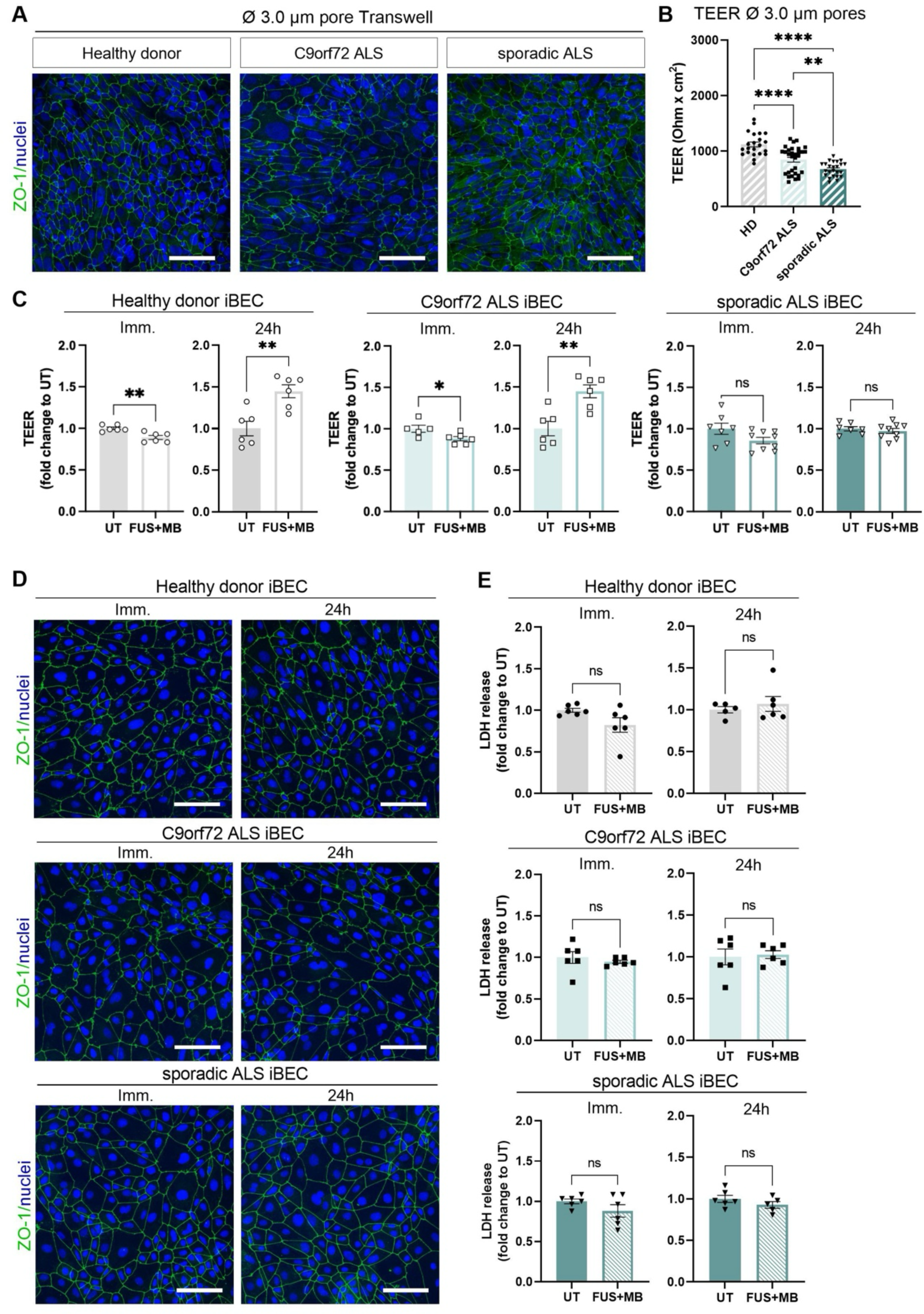
Effects of FUS^+MB^ on viability and monolayer integrity of control and ALS iBECs. **(A)** Representative immunofluorescence images ZO-1 (green) with Hoechst nuclear counterstaining (blue) in healthy donor, C9orf72 ALS, and sporadic ALS iBECs cultured in 3.0 µm pore Transwell insert membrane. Scale bar = 100 µm. (**B**) Trans-endothelial electrical resistance (TEER, Ohm x cm^2^) of healthy donor, C9orf72 ALS and sporadic ALS iBECs, measured in 3.0 µm pore Transwells. A minimum of *n* = 22 independent replicates per line. (**C**) Changes in TEER of healthy donor, C9orf72 ALS, and sporadic ALS iBECs exposed to FUS^+MB^, immediately and 24 h following the treatment. TEER was measured in 3.0 µm pore Transwells and shown as fold changes to untreated (UT) cells at respective time points. A minimum of *n* = 5 independent replicates per line. (**D**) Representative immunofluorescence images ZO-1 (green) with Hoechst nuclear counterstaining (blue) in healthy donor, C9orf72 ALS, and sporadic ALS iBECs exposed to FUS^+MB^, shown immediately and 24 h following the treatment. Scale bar = 100 µm. (**E**) Changes in relative lactate dehydrogenase (LDH) release in healthy donor, C9orf72 ALS, and sporadic ALS iBECs exposed to FUS^+MB^, immediately and 24 h following the treatment. LDH release shown as fold changes to respective untreated (UT) cells at each time point. A minimum of *n* = 5 independent replicates per line. Data in (B) analysed with One-way ANOVA with Tukey’s test, and in (C, E) with Student’s *t*-test or Welch’s *t*-test. Error bars = SEM. \**P*<0.05, \*\**P*<0.01, \*\*\**P*<0.001, \*\*\*\**P*<0.0001. Imm.- immediately.

Utilising the 3.0 µm pore model, we first examined the effects of FUS^+MB^ on iBECs barrier integrity. In clinical and animal studies FUS^+MB^ was shown to induce rapid BBB opening followed by full BBB integrity recovery within 24 h (28,32,53,59). Analogous to *in vivo* effects (28,32,53,59), we found a significant decrease in TEER of healthy donor (*P*<0.01) and C9orf72 ALS (*P*<0.05) iBECs immediately following FUS^+MB^ treatment (Figure 3C). While achieving a similar fold reduction in TEER in healthy donor and C9orf72 ALS iBECs, statistically only a trend towards decreased TEER was found in sporadic ALS iBECs (healthy donor: 0.8905 ± 0.02, C9orf72 ALS: 0.8803 ± 0.03, sporadic ALS: 0.8565 ± 0.04, mean ± SEM fold change vs respective untreated control, Figure 3C, **Figure S4A**). This was also consistent with the estimated 10% reduction in TEER induced by 0.3 MPa FUS^+MB^ in other *in vitro* BBB studies (37,60). At 24 h following the treatment, no difference was found in TEER of sporadic ALS iBECs, while TEER was increased (*P*<0.01) in healthy donor and C9orf72 ALS iBECs (Figure 3C, **Figure S4A**), corresponding to our respective observations in sporadic (37) and familial (34) AD models. To confirm that changes in TEER were not resulting from cell death induced by FUS^+MB^ we next assessed iBECs viability immediately and 24 h following FUS^+MB^ exposure. Our analysis revealed that FUS^+MB^ treatment did not cause any visible damage in the control and ALS iBECs monolayer (Figure 3D) and had no adverse effects on cell viability as assessed by lactate dehydrogenase (LDH) release assay at either time point tested (Figure 3E). Relative LDH release also did not differ between the control and ALS lines immediately and 24 h post FUS^+MB^ (**Figure S4B**).

### FUS^+MB^elicits differential molecular bioeffects in control and ALS iBECs

Although ultrasound-based ALS BBB opening technology is rapidly moving towards clinical use (32), the molecular responses of human ALS BBB cells to FUS^+MB^ remain unexplored. To further investigate the effects of FUS^+MB^ on control and ALS iBECs, we exposed the cells to FUS^+MB^ and evaluated the resulting changes in gene expression at two time points corresponding to iBEC monolayer opening and its recovery. Since FUS^+MB^ was previously shown to elicit molecular changes in tight and adherens junctions (34,61) we first examined the expression of cell junction markers in iBECs via two-step reverse transcription quantitative polymerase chain reaction (RT-qPCR). When comparing to respective untreated cells, we found a significant decrease (*P*<0.01) in *OCLN* and *CDH5* expression in healthy donor iBECs immediately post FUS^+MB^ while *OCLN* and *CLDN5* demonstrated reduced, (*P*<0.05) and *TJP1* increased, (*P*<0.05) expression in C9orf72 ALS iBECs (Figure 4A, **Figure S5**). Interestingly, we found no changes in tight and adherens junction gene expression levels in sporadic ALS iBECs immediately following FUS^+MB^ when compared to untreated cells from this line (Figure 4A, **Figure S5**). Similarly, the expression of drug transporter *ABCB1* was not altered immediately post FUS^+MB^ in healthy donor and ALS iBECs (Figure 4A, **Figure S5**). However, when comparing between examined iBEC lines, we found differences in *OCLN*, *CLDN5, TJP1* and *CDH5* expression immediately after FUS^+MB^ treatment collectively suggesting the molecular bioeffects of FUS^+MB^ may potentially impact different pathways in control and ALS cells (Figure 4B).

**Figure 4.**
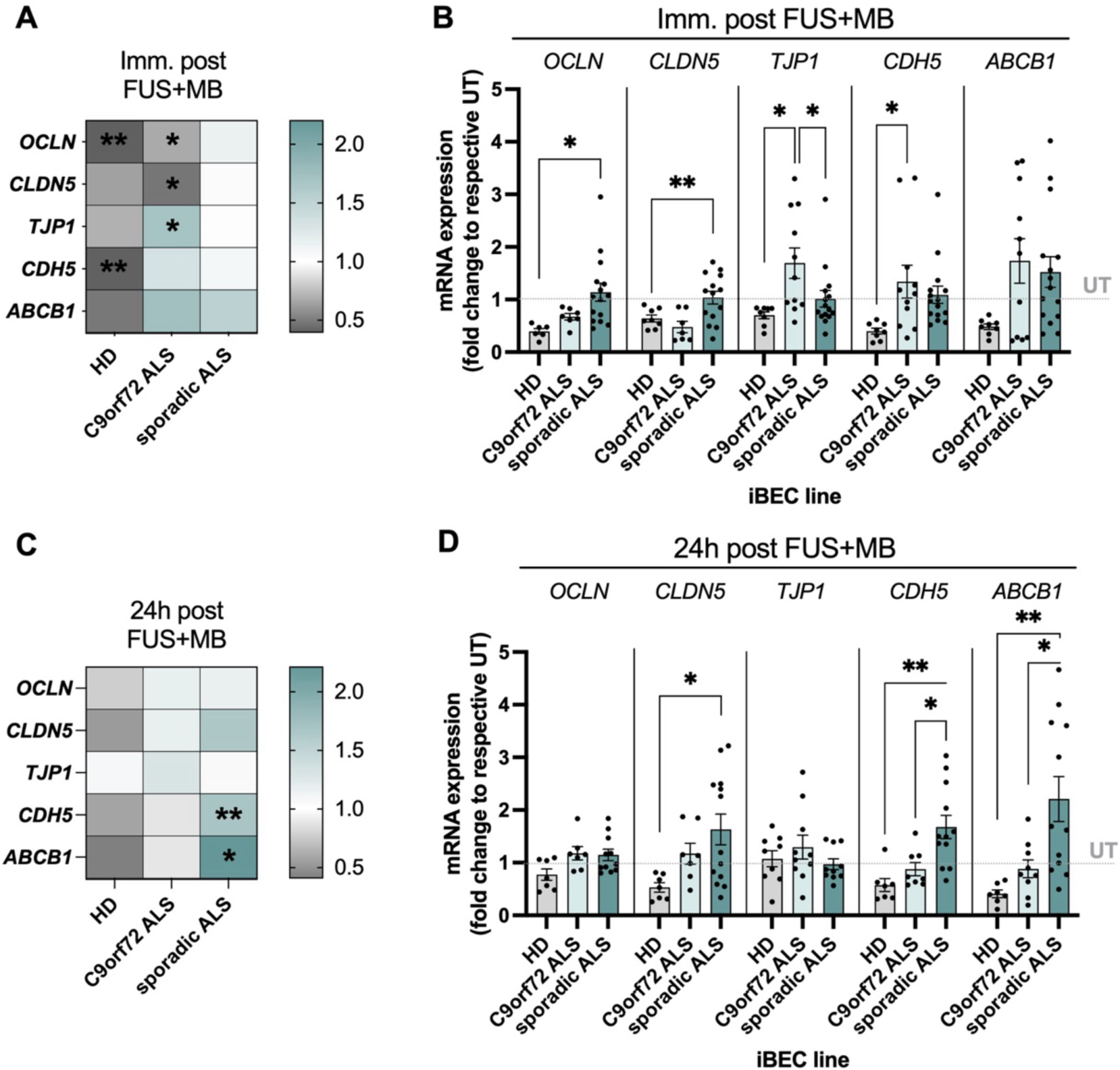
Effects of FUS^+MB^ on the expression of junctional and transporter marker genes in control and ALS iBECs. (**A**,**C**) Heatmap summarising fold changes changes in mRNA expression of *OCLN*, *CLDN5, TJP1*, *CDH5*, *ABCB1* in healthy donor, C9orf72 ALS, and sporadic ALS iBECs exposed to FUS^+MB^, immediately (A) and 24 h (C) following the treatment. (**B**,**D**) Comparison of changes in mRNA expression of *OCLN*, *CLDN5, TJP1*, *CDH5*, *ABCB1* between healthy donor, C9orf72 ALS, and sporadic ALS iBEC, immediately (B) and 24 h (D) following the FUS^+MB^ treatment. In A-D, a minimum of *n* = 6 independent replicates per line. Data showed as fold changes to respective untreated (UT) cells from each cell group at each time point. Data in (A, C) analysed with Student’s *t*-test or Welch’s *t*-test and data in (B, D) analysed with One-way ANOVA with Tukey’s test, error bars = SEM. Only significant changes are displayed in A-D. \**P*<0.05, \*\**P*<0.01. Imm.-immediately.

24 h post FUS^+MB^, tight and adherens junction gene expression normalised in healthy donor and C9orf72 ALS iBECs (Figure 4C, **Figure S5**), suggesting iBEC monolayer recovery in these lines. Interestingly, at this time point, we found a significant increase in *CDH5* (*P*<0.01) and *ABCB1* (*P*<0.05) expression in sporadic ALS iBECs when compared to respective untreated cells (Figure 4C, **Figure S5**). At 24 h, FUS^+MB^-treated ALS iBECs also demonstrated increased expression of *CLDN5* as compared to healthy donor iBECs and higher expression of *CDH5* and *ABCB1* when compared to healthy donor and C9orf72 ALS cells (Figure 4D), together indicating temporarily distinct responses of sporadic ALS iBECs to FUS^+MB^.

FUS^+MB^-mediated BBB opening was shown to remodel the immune landscape in the murine brain (62–66). However, the neuroinflammatory responses to FUS^+MB^ of human ALS BBB cells are currently unknown. Thus, as both human primary BEC and iBECs have shown the capacity to elicit inflammatory responses (35,67,68), we next investigated the expression of inflammatory mediators in FUS^+MB^-treated iBEC. In this exploratory analysis inflammatory markers interleukin-6 (*IL6*), interleukin-8 (*IL8*), interleukin-1A (*IL1A*), C-C motif chemokine ligand 2 (*CCL2*) and tumor necrosis factor-α (*TNF*) were selected as they were previously implicated in immune responses to FUS^+MB^ (37,65,66) and were shown to be expressed by iBECs (35).

Since neuroinflammation was linked with BBB pathology in ALS (21,69), we first examined the baseline expression of inflammatory marker genes in untreated healthy donor and ALS iBECs. Interstingly, while ALS is commonly associated with an increased proinflammatory profile (1,23,39,70), we found a decreased (*P*<0.05) expression of proinflammatory marker genes, *IL6* and *TNF,* in sporadic ALS iBECs compared to healthy donor cells, with *IL6* demonstrating also reduced (*P*<0.0001) expression in sporadic ALS iBECs compared to C9orf72 ALS iBECs (Figure 5A).

**Figure 5.**
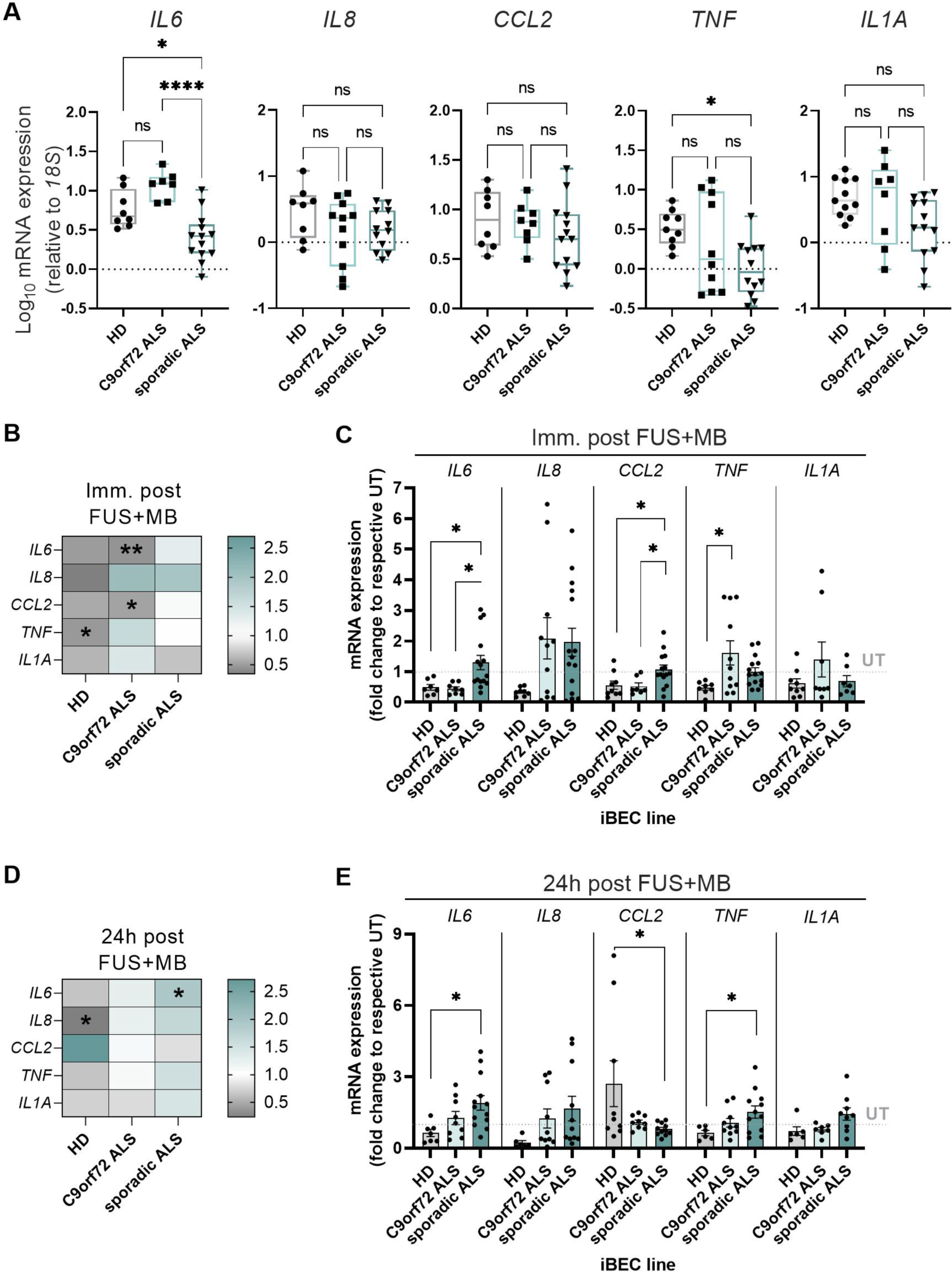
Effects of FUS^+MB^ on the expression of inflammatory marker genes in control and ALS iBECs. (**A**) Relative mRNA expression of inflammatory marker genes *IL6*, *IL8, CCL2*, *TNF* and *IL1A* in the untreated healthy donor, C9orf72 ALS, and sporadic ALS iBECs. Data presented as Log10 of ΔΔCT x 10^6^, relative to *18S*. (**B**,**D**) Heatmap summarising fold changes in mRNA expression of *IL6*, *IL8, CCL2*, *TNF* and *IL1A* in healthy donor, C9orf72 ALS, and sporadic ALS iBEC exposed to FUS^+MB^, immediately (B) and 24 h (D) following the treatment. (**C**,**E**) Comparison of changes in mRNA expression of *IL6*, *IL8, CCL2*, *TNF* and *IL1A* between healthy donor, C9orf72 ALS, and sporadic ALS iBECs, immediately (C) and 24 h (E) following the FUS^+MB^ treatment. In A-E, a minimum of *n* = 6 independent replicates per line. Data in (B-E) showed as fold changes to respective untreated (UT) cells from each cell group at each time point. Data in (B, D) analysed with Student’s *t*-test or Welch’s *t*-test and data in (A, C, E) analysed with One-way ANOVA with Tukey’s test, error bars = SEM. Only significant changes are displayed in B-E. \**P*<0.05, \*\**P*<0.01, \*\*\*\**P*<0.0001. Imm.-immediately.

When assessing gene expression changes immediately following FUS^+MB^, we identified decreased expression of *TNF* (*P*<0.05) in healthy donor cells as well as a decrease in *IL6* (*P*<0.01) and *CCL2* (*P*<0.05) in C9orf72 ALS iBECs suggesting potential immunosuppressive effect of FUS^+MB^ in these lines (Figure 5B, **Figure S6**). Similar to the junctional markers, we found no changes in inflammatory marker gene expression in sporadic ALS iBECs immediately post FUS^+MB^ (Figure 5B, **Figure S6**). At 24 h, expression of *IL8* was reduced (*P*<0.05) in healthy donor iBECs while *IL6* showed increased (*P*<0.05) expression in sporadic ALS iBECs (Figure 5D, **Figure S6**). When immediate responses were compared between independent lines, we found expression of *IL6* and *CCL2* to be decreased (*P*<0.05) in healthy donor and C9orf72 ALS iBECs compared to sporadic ALS iBECs while *TNF* showed increased (*P*<0.05) expression in C9orf72 iBECs compared to healthy donor control (Figure 5C). Following 24 h, *IL6* and *TNF* expression were increased (*P*<0.05) in sporadic ALS iBECs compared to healthy donor cells and *CCL2* expression was reduced (*P*<0.05). Together, this differential gene expression may suggest that iBECs from each individual donor generate a unique and temporarily dynamic inflammatory profile in response to FUS^+MB^.

### FUS^+MB^ enhances anti-TDP-43 antibody permeability at ALS BBB in vitro

Different ALS drugs were successfully delivered across the BBB with FUS^+MB^ in the animal studies (29–31). Yet, improving ALS drug delivery with FUS^+MB^ has not been trialled in the human ALS BBB *in vitro* or ALS patients. Here we focused on testing the delivery of a potentially therapeutic antibody in our ALS BBB model given the previous broad preclinical success of antibody and FUS^+MB^ combined therapy (37,41,71–73) indicating high compatibility of ultrasound technique with this drug format. Due to the lack of immunotherapeutic agents approved for ALS, we selected a commercially available anti-TDP-43 antibody as a model molecule (∼150 kDa) and developed a platform to trial its delivery with FUS^+MB^ in ALS iBECs (Figure 6A).

**Figure 6.**
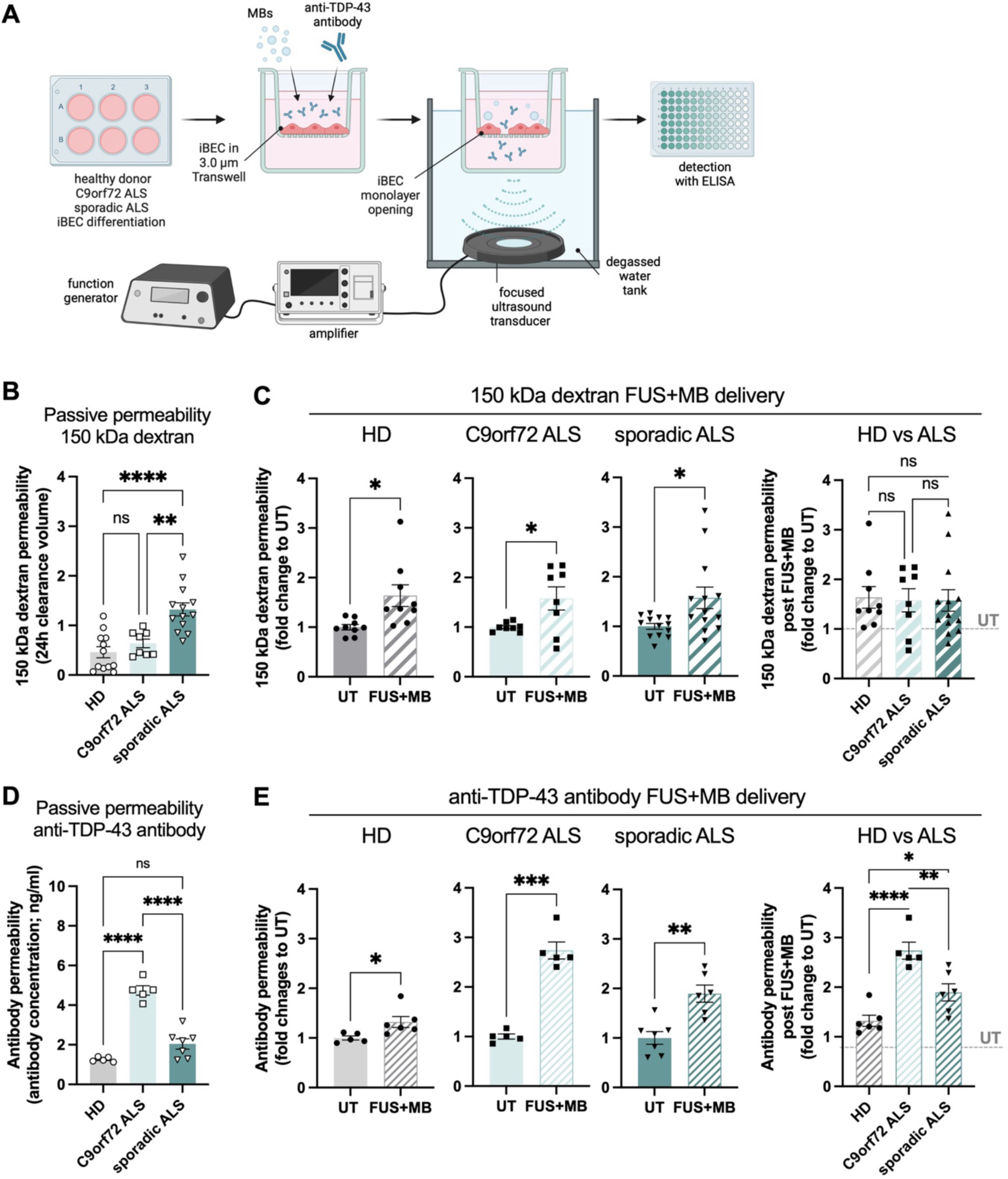
FUS^+MB^-mediated delivery of 150 kDa dextran and anti-TDP-43 antibody in control and ALS iBECs. (**A**) Schematic illustration of FUS^+MB^ mediated anti-TDP-43 antibody delivery in the Transwell model. (**B**) Passive permeability of 150 kDa dextran in healthy donor, C9orf72 ALS, and sporadic ALS iBECs cultured in 3.0 µm poreTranswell. A minimum of *n* = 8 independent replicates per line. Data presented as clearance volume of dextran at 24 h. (**C**) Delivery of 150 kDa dextran in healthy donor, C9orf72 ALS, and sporadic ALS iBECs using FUS^+MB^. Permeability shown as fold change to respective untrated (UT) cells at 24 h. A minimum of *n* = 8 independent replicates per line. (**D**) Passive permeability of anti-TDP-43 antibody in healthy donor, C9orf72 ALS, and sporadic ALS iBECs cultured in 3.0 µm pore Transwell. A minimum of *n* = 5 independent replicates per line. Data presented as anti-TDP-43 IgG concentration (ng/ml) detected in the bottom of the Transwell at 24 h. (**E**) Delivery of anti-TDP-43 antibody in healthy donor, C9orf72 ALS, and sporadic ALS iBECs using FUS^+MB^. Permeability shown as fold change to respective untrated (UT) cells at 24 h. A minimum of *n* = 5 independent replicates per line. Data in (B, D) and three-group comparison in (C,E) analysed with with One-way ANOVA with Tukey’s test. Two-group comparisons in (C,E) analysed with Student’s *t*-test or Welch’s *t*-test. \**P*<0.05, \*\**P*<0.01, \*\*\**P*<0.001, \*\*\*\**P*<0.0001, error bars = SEM.

First, the passive transport of large molecules in ALS iBECs was explored by studying the permeability of a biologically inert 150 kDa dextran fluorescent tracer. Similar to the 5 kDa tracer (Figure 1C), we found increased leakage of 150 kDa dextran in sporadic ALS iBECs compared to healthy donor (*P*<0.0001) and C9orf72 ALS (*P*<0.01) iBECs (Figure 6B). This suggests that the sporadic ALS cells demonstrate iBECs permeability impairment. Next, cells were cultured on 3.0 µm pore Transwells and exposed to FUS^+MB^ together with 150 kDa dextran and its permeability was assessed at 24 h post-treatment as we previously described (37). Similar to our observations in the AD model (37), we found FUS^+MB^ to increase (*P*<0.05) 150 kDa dextran delivery in iBECs with no differences in delivery efficiency between control and ALS lines (Figure 6C).

Interestingly, when the passive permeability of anti-TDP-43 antibody was assessed, we identified its increased (*P*<0.0001) passive permeability in C9orf72 ALS iBECs, compared to healthy donor and sporadic ALS iBECs (Figure 6D), with this effect not directly corresponding to C9orf72 ALS iBEC monolayer TEER (Figure 3B) or permeability to 150 kDa dextran (Figure 6B). We next trialled anti-TDP-43 antibody delivery with FUS^+MB^ and found its increased permeability in healthy donor (*P*<0.05), C9orf72 ALS (*P*<0.001) and sporadic ALS (*P*<0.01) iBECs (Figure 6E). Analogous to passive permeability, FUS^+MB^-enhanced anti-TDP-43 antibody delivery efficiency proved to be the highest in C9orf72 ALS iBECs (*P*<0.0001 vs HD iBECs and *P*<0.01 vs sporadic ALS iBECs, Figure 6E). Importantly, a higher fold permeability increase (*P*<0.05) was also found in sporadic ALS iBECs compared to healthy donor cells (Figure 6E), collectively suggesting the high efficiency of anti-TDP-43 antibody delivery with FUS^+MB^ in ALS models. Furthermore, despite BBB integrity changes not being detectable with TEER technique in sporadic ALS iBECs (Figure 3C), observation of increased permeability of 150 kDa dextran and anti-TDP-43 antibody confirmed effective monolayer opening in ALS iBECs following FUS^+MB^.

## DISCUSSION

ALS is a fatal neurodegenerative condition with approved treatment options being limited to riluzole (74), edaravone (75) and tofersen antisense oligonucleotide (10,76) therapies. Their effectiveness however is restricted by the pharmacoresistance of the BBB (10,76–79) warranting the development of non-invasive drug delivery technologies that could maximise the amount of drug reaching its CNS target. FUS^+MB^ is a promising technique facilitating temporary BBB opening (28) with its clinical safety and effectiveness in ALS confirmed by a pioneering study (32). While reversibly increasing the permeabilisation of the BBB to applied drugs in animal studies (29–31), FUS^+MB^ has not yet been combined with any therapeutic molecule in ALS patients.

Here we developed an ALS patient-cell-derived *in vitro* platform to screen for the molecular and cellular bioeffects of FUS^+MB^ and provide the first proof-of-concept evidence for the feasibility of FUS^+MB^ to enhance large molecule drug delivery at human ALS BBB. By utilising sporadic and familial (*C9orf72* expansion variant) ALS iBECs as a BBB model, we found phenotypical differences in ALS iBECs integrity, permeability, intracellular TDP-43 protein localisation, junctional marker and drug transporter expression, replicating some of the hallmark disease-associated changes at BBB in ALS (15,16). Interestingly, familial ALS cells were obtained from asymptomatic (at the time of skin biopsy) *C9orf72* expansion carrier who experienced disease onset later in life, suggesting these BBB changes may precede clinical ALS symptoms development in *C9orf72*-dependent ALS.

Our exploratory analysis further revealed that FUS^+MB^ elicits temporary changes in ALS iBEC monolayer integrity and permeability to fluorescent 150 kDa tracer with no adverse effects on cell viability or morphology. We were also the first to demonstrate improved transport of anti-TDP-43 antibody in ALS iBECs following FUS^+MB^ exposure. These effects were accompanied by molecular changes in junctional protein, drug transporter and inflammatory marker gene expression in ALS iBECs providing important insights on the cell-specific responses of human BBB cells to FUS^+MB^. With our model recapitulating aspects of clinical BBB phenotype in ALS (15,16,24), we believe observed effects would prove to be of high translational relevance and support future clinical success of FUS^+MB^ mediated drug delivery in ALS patients.

Extensive evidence suggests BBB impairment in ALS (15,16,19,26,52). However, with important discrepancies found in BBB pathology between ALS animal models and patients (24), it remains critical to develop fully human *in vitro* BBB models facilitating understanding of BBB changes in ALS. iPSC-derived iBECs (38,43,54) serve as key cellular component of human BBB models, offering currently the highest drug permeability mimicry when compared to the human *in vivo* brain (80,81). Despite the importance of BBB modelling in ALS, studies describing iBECs generation from ALS patient iPSCs are sparse and limited to its familial form (58). Our phenotypic characterisation of human ALS iBECs revealed a decrease in TEER of monolayers formed by ALS cells, suggesting barrier integrity impairment in the disease, previously also reported for SOD1 mutation and *C9orf72* expansion carrying iBECs (58). We have also found the barrier integrity to be lower in sporadic ALS iBECs as compared to C9orf72 iBECs and accompanied by increased passive permeability to 5 kDa and 150 kDa fluorescent tracers, pointing to potential phenotypic heterogeneity between disease subtypes. This could also explain inconsistent results of clinical reports where BBB disruption was found only in a fraction of ALS patients (82–85), highlighting the need for more careful patient stratification in generalised cohort studies.

Interestingly, we also found sporadic ALS iBECs to express lower levels of *CDH5* encoding for adherens junction complex protein VE-cadherin, previously shown to govern endothelium permeability (86). With no changes observed in tight junction-associated genes, these results could indicate specific vulnerability of adherens junction complexes in sporadic ALS. Furthermore, in contrast to previous *in vivo* studies (13,48), experiments performed in our model identified reduced expression of *ABCB1* in sporadic ALS iBECs with its expression not being altered in C9orf72 ALS iBECs. This however was partially aligned with an *in vitro* study where P-pg activity was shown to be decreased in *C9orf72* expansion carrying iBECs (58), together supporting overall P-gp dysregulation in ALS.

TDP-43 protein pathology can be found in almost all ALS cases where it is primarily linked to motor neuron degeneration (87,88). Interestingly, TDP-43 and phosphorylated TDP-43 aggregates were recently found in cortical vessel walls of sporadic ALS patients (52). However, with BBB comprising multiple cell types, an association of TDP-43 with specific cell type within the neurovascular unit was not determined (52). Similarly, the source (endogenous vs neuron-derived) of TDP-43 protein aggregates at the blood vessels was not examined and the brain endothelial cell-type-specific effects of TDP-43 proteinopathy are not clear.

Here we found cytoplasmic TDP-43 protein aggregates in a single-cell-type ALS iBECs culture. This was accompanied by the disease-associated changes in ALS iBECs with the degree of TEER and permeability impairment being proportional to the amount of TDP-43 aggregates found in C9orf72 and sporadic ALS cells. Although further investigation is required, our observation suggests that endogenous TDP-43 proteinopathy may be present in BECs and influence cerebrovascular homeostasis in the disease. Importantly, however, iPSCs utilised in this study were reprogrammed from skin fibroblasts of a healthy donor (34) and ALS patients, and cytoplasmic mislocalisation and accumulation of TDP-43 was recently reported in human ALS fibroblasts (89–91). Hence TDP-43 pathology could also be carried from source fibroblasts, warranting further validation of our *in vitro* results in human ALS brain tissue.

When assessing the passive transport of molecules in our ALS model we found increased permeability of biologically inert 5 kDa and 150 kDa dextran in sporadic ALS iBECs, consistent with the lowest TEER generated by this line. However, anti-TDP-43 antibody leakage was the highest in C9orf72 ALS cells, with C9orf72 ALS iBECs demonstrating higher barrier integrity and simultaneously lower cytoplasmic TPD-43 protein content than sporadic ALS iBECs. This may suggest that the permeability of potentially therapeutic antibodies in ALS iBECs is not dependent on the net barrier integrity, but rather, points to the complex molecular interactions of antibodies with patient-derived cells; mimicking our findings for anti-tau antibodies in sporadic AD model (41). Observed effects may also imply the heterogeneity in drug permeability at barriers of sporadic and familial ALS patient subtypes and highlight alterations in paracellular and transcellular BBB transport pathways in ALS. This could also explain the lack of widespread drug permeability at seemingly more leaky BBB in ALS (92) since all three approved treatments (13,76) require at least in part transcellular transport mechanisms.

Given the need for improving drug delivery at the ALS BBB (92) and the lack of fully human *in vitro* models allowing for investigation of FUS^+MB^ effects on ALS BBB cells, we next utilised characterised ALS iBECs to develop a FUS^+MB^-mediated drug permeability screening platform. Correlating with clinical ALS study (32), our analysis confirmed the transient effect of FUS^+MB^ on iBEC monolayer opening, as reflected by a reduction in TEER immediately following the treatment. While no changes were found in the integrity of sporadic ALS iBECs 24 h post-treatment, the TEER of healthy donor control and familial C9orf72 ALS iBECs was increased as compared to untreated cells at this time point. Interestingly, in our previous studies with sporadic (37) and familial (34) AD models, we also found a consistent decrease in barrier integrity immediately following FUS^+MB^; whereas the TEER was no longer altered at 24 h post FUS^+MB^ in the sporadic AD model, but increased in familial AD iBECs, mirroring respective observations in sporadic and familial ALS cells. Together this could suggest some cellular responses to FUS^+MB^ could be shared between familial and sporadic variants of neurodegenerative diseases warranting further investigation.

We have previously performed RNA sequencing in sporadic AD iBECs exposed to FUS^+MB^ and reported no differences in gene expression immediately and 24 h post-treatment (37). Our other analysis conducted by qPCR demonstrated FUS^+MB^-induced changes in *OCLN*, *CLDN5*, *TJP1* and *CDH5* expression in familial AD iBECs (34). In this study, when considering FUS^+MB^ elicited molecular effects in ALS cells, we found expression of the same junctional marker genes to be altered in iBECs. However, this was limited to healthy donor and C9orf72 cells, with sporadic ALS iBECs showing no response to FUS^+MB^ immediately following the treatment. Sporadic ALS iBECs proved to be more responsive at 24 h timepoint presenting altered *CDH5* expression levels. Our analysis also revealed a trend towards decreased expression of P-gp-encoding *ABCB1* in healthy donor cells and contrarily increased expression in sporadic ALS iBECs following FUS^+MB^, supporting our previously reported modulatory effects of FUS^+MB^ on drug transporter expression and function in AD iBECs (93). With P-pg serving as a major drug efflux transporter at the BBB, its elevated activity could prove to be counteractive to FUS^+MB^-mediated drug delivery and confer CNS pharmacoresistance in sporadic ALS. While requiring validation at protein and functional levels, further investigation in our ALS model could aid in the early identification of such responses and test for strategies (13) mitigating undesired effects of FUS^+MB^ in defined subpopulations of patients.

FUS^+MB^ was previously shown to induce neuroinflammatory responses in animal studies (65,66,94) motivating our exploratory analysis of proinflammatory marker expression in ALS iBECs. Intriguingly, we identified reduced expression of *TNF* and *IL8* in healthy donor cells, and *IL6* and *CCL2* in C9orf72 ALS iBECs, suggesting FUS^+MB^ may rather confer immunosuppressive effects in cells of some individuals. Sporadic ALS iBECs showed a differential expression profile, presenting the increased expression of *IL6* and *CCL2* at the immediate time point and *IL6* and *TNF* at 24 h when compared to other lines, but overall lacked a profound proinflammatory response to FUS^+MB^. These observations were consistent with our previous study (37) showing no change or decreased proinflammatory marker gene expression in sporadic AD iPSC-derived iAstrocytes exposed to FUS^+MB^. While *in vivo* responses in the human brain are still unexplored, this could point to potential discrepancies in neuroinflammatory effects of FUS^+MB^ between animal and human cells highlighting the importance of studies in fully human *in vitro* models.

With BBB limiting an estimated 99.8 % of peripherally administered large molecule drugs such as therapeutics antibodies (95), we next tested the possibility of enhancing the permeabilisation of ALS BBB to large molecules with FUS^+MB^. As a proof-of-concept, we trialled the delivery of anti-TDP-43 antibody, considering promising preclinical development of TDP-43 targeted immunotherapies (96–99) as well as the recent clinical success of therapeutic antibodies in the treatment of other neurodegenerative disorders (100–104). Although the utilised anti-TDP-43 antibody has not yet proved therapeutically beneficial, we were previously successful in delivering

Aducanumab (Aduhelm^TM^) and anti-tau therapeutic antibodies with FUS^+MB^ in human Alzheimer’s disease *in vitro* models (37,41); and a recent clinical study demonstrated beneficial effects of Aducanumab combined with FUS^+MB^ in Alzheimer’s disease patients, as compared to Aducanumab alone (104). Hence, we hypothesised that analogously, achieving increased permeability of this model anti-TDP-43 antibody in our cell platform could be representative of future clinically approved ALS immunotherapeutics.

Here, we were able to demonstrate for the first time that FUS^+MB^ facilitates improved permeability of anti-TDP-43 antibody at ALS BBB *in vitro*. Interestingly, antibody delivery efficiency was higher in C9orf72 and sporadic ALS iBECs than control cells, reaching 2.7-and 1.9-fold increases respectively, as compared to its passive permeability, suggesting potential for promising drug delivery enhancement in ALS. While improving the cost-effectiveness of immunotherapy, it remains to be determined whether such improvement in drug delivery would be therapeutically meaningful in patients. However, a 2-fold increase in edaravone permeability following FUS^+MB^ was associated with additional symptom amelioration in an ALS mouse model (30), suggesting it is a sufficient increase to enhance therapeutic outcomes. In the future, further increase in drug permeability can be achieved in combination with currently preclinically investigated tight junction binders which were shown to improve BBB opening with FUS^+MB^, expanding the therapeutic window for drug application (60).

While providing an advancement in FUS^+MB^ drug delivery in ALS, there are several limitations to this study. Firstly, our observations should be confirmed with an increased number of iPSC lines (105) to account for broader heterogeneity in sporadic and familial ALS patient populations. The development of multicellular and 3D models (37,106) would also be essential for a complete understanding of FUS^+MB^ effects on the neurovascular unit in ALS. Finally, the BBB is not the only barrier limiting drug delivery into the CNS and the blood-spinal cord barrier (BSCB) is also an important contributor to pharmacoresistance in ALS. Although FUS^+MB^-mediated BSCB opening and drug delivery into the spinal cord have been successfully performed in non-ALS rodents (107–109), this technique has not yet been used clinically in humans. Therefore, in the current study, we have focused on the development of the BBB model for FUS^+MB^ mediated drug screening in

ALS as this technology can be more rapidly delivered to patients. However, future *in vitro* BSCB modelling and FUS^+MB^-mediated drug delivery screening would be a necessary advancement in overcoming CNS barriers in ALS.

Together these results provide important proof-of-concept evidence for non-invasive, FUS^+MB^-facilitated antibody transport across the *in vitro* ALS BBB, which could aid in improving the delivery of large molecule drugs and TDP-43-directed immunotherapies (96–99) in ALS. With no such platform reported to date, our study provides the first patient-cell-derived model for investigation of FUS^+MB^ bioeffects and screening of FUS^+MB^-deliverable drug formats most compatible with disease subtypes, supporting personalised medicine approaches to ALS drug delivery.

## CONCLUSIONS

The BBB restricts the entry of large-molecule drugs into the brain, limiting treatment effectiveness in ALS. Our study presents a novel ALS patient-derived blood-brain barrier (BBB) cell platform for the advancement of FUS^+MB^-mediated drug delivery and demonstrates for the first time that FUS^+MB^ increases the delivery of potentially therapeutic anti-TDP-43 antibody across the *in vitro* barrier. These findings are an important step in accelerating the translation of FUS^+MB^ as a CNS drug delivery technology in the treatment of ALS, especially in conjunction with immunotherapeutics.

## Abbreviations

ALS: Amyotrophic lateral sclerosis
AD: Alzheimer’s disease
BBB: blood-brain barrier
BEC: brain endothelial cell
BSCB: blood-spinal cord barrier
CNS: central nervous system
ELISA: enzyme-linked immunosorbent assay
FUS: focused ultrasound
HD: healthy donor
iBEC: induced brain endothelial-like cell
Imm: immediately
iPSC: induced pluripotent stem cell
MB: microbubble
qPCR: quantitative polymerase chain reaction
TDP-43: TAR DNA-binding protein 43
TEER: trans-endothelial electrical resistance
UT: untreated

## DECLARATIONS

### Ethics approval

The study was approved by the University of Palermo Ethics Committee (Palermo 1, n° 4/2019), the University of Wollongong Human Research Ethics Committee (18/366) and QIMR Berghofer Human Ethics Committee (P2197). All participants provided informed consent before participating in the study.

### Consent for publication

Not applicable.

### Availability of data and materials

The datasets supporting the conclusions of this article are included within the article and its additional file and are available from the corresponding author upon reasonable request.

## CRediT authorship contribution statement

**JMW**: conceptualisation, methodology, investigation, formal analysis, visualisation, writing - original draft, writing - review & editing; **MCC**: methodology; **MP**: methodology; **THN**: methodology; **VLB**: conceptualisation, methodology, resources; **LEO**: methodology; **LO**: conceptualisation, methodology, resources; **ARW**: conceptualisation, writing - review & editing, supervision, project administration, funding acquisition. All authors reviewed and approved the final manuscript.

## Acknowledgements

We thank QIMR Berghofer MRI Flow Cytometry and Imaging Facility for their assistance and Dr Satomi Okano and QIMR Berghofer MRI Statistics department for their advice on data analysis. We thank prof. Jose M. Polo (Monash University) for the provision of HDFa hiPSC line. Graphical elements were generated with Biorender.com.

