## Supplemental material for "A patient-derived amyotrophic lateral sclerosis blood-brain barrier cell model reveals focused ultrasound-mediated anti-TDP-43 antibody delivery"

**SUPPLEMENTAL FIGURES:**

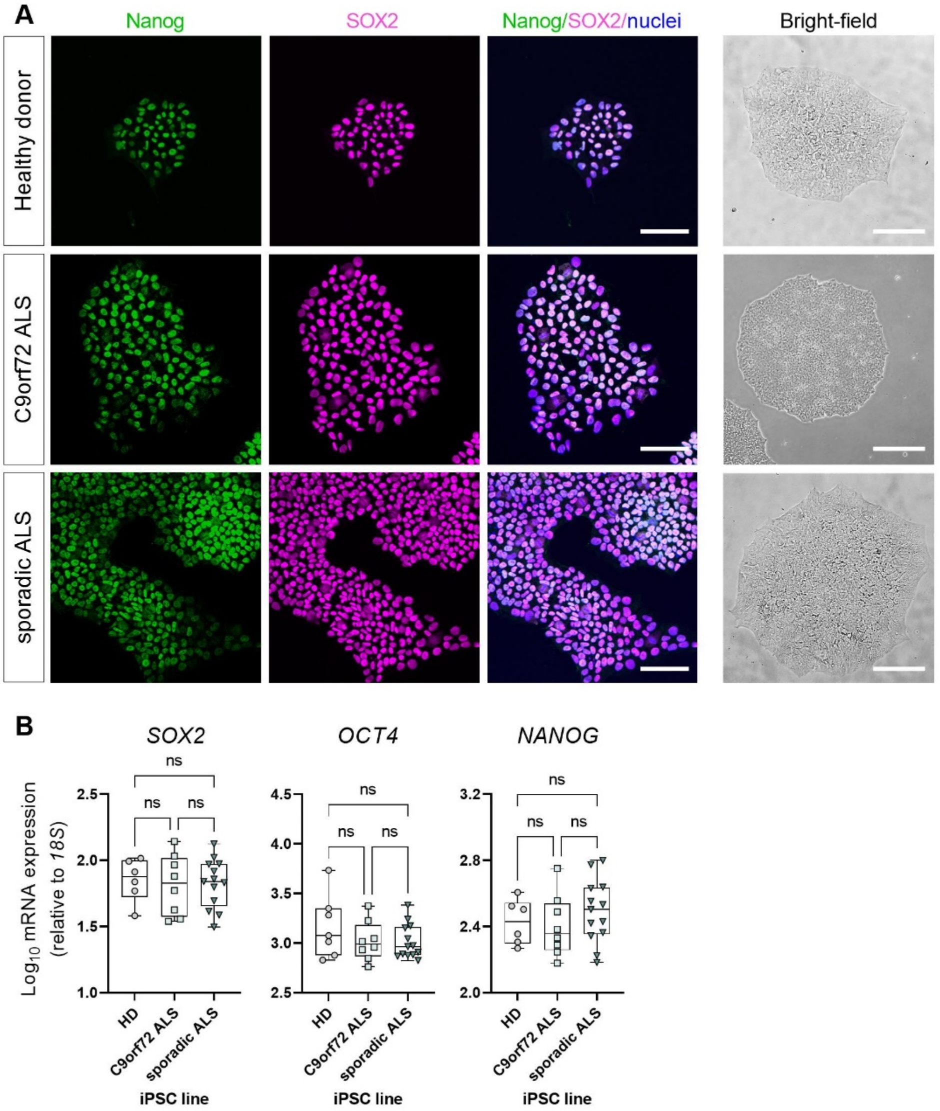

**Supplementary figure S1. Characterisation of healthy donor and ALS human induced pluripotent stem cells (iPSC).** (**A**) Representative immunofluorescence images of Nanog (green) and SOX2 (magenta) with Hoechst nuclear counterstaining (blue) and bright-field image of healthy donor, C9orf72 ALS, and sporadic ALS iPSC. Scale bar = 100 µm in immunofluorescence images and 200 µm in brightfield images. (**B**) Relative mRNA expression of pluripotency marker genes *SOX2*, *OCT4* and *NANOG* in healthy donor, C9orf72 ALS, and sporadic ALS iPSC. Data presented as $\text{Log}_{10}$ of $\Delta\Delta$CT x $10^6$, relative to *18S*. A minimum of $n$ = 6 independent replicates per line. Data in (B) analysed with One-way ANOVA with Tukey's test, error bars = SEM.

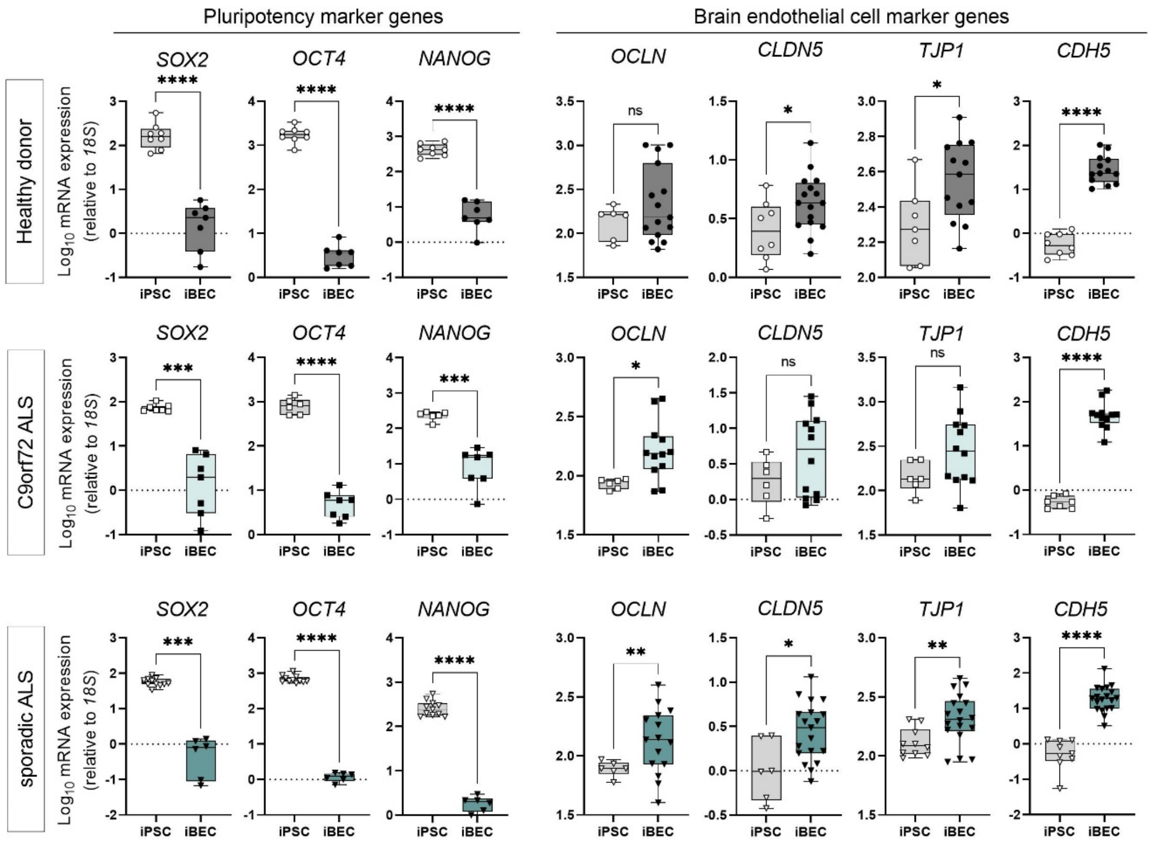

**Supplementary figure S2. Relative mRNA expression of pluripotency and brain endothelial cell markers in control and ALS iPSCs and iBECs.** Relative mRNA expression of pluripotency marker genes *SOX2*, *OCT4* and *NANOG* and brain endothelial cell marker genes *OCLN*, *CLDN5, TJP1* and *CDH5* in healthy donor, C9orf72 ALS, and sporadic ALS iPSCs and iBECs. Data presented as $Log_{10}$ of ΔΔCT x $10^6$, relative to *18S*. A minimum of $n$ = 5 independent replicates per line. Data analysed with Student's *t*-test or Welch's *t*-test. **P*<0.05, ***P*<0.01, ****P*<0.001, *****P*<0.0001.

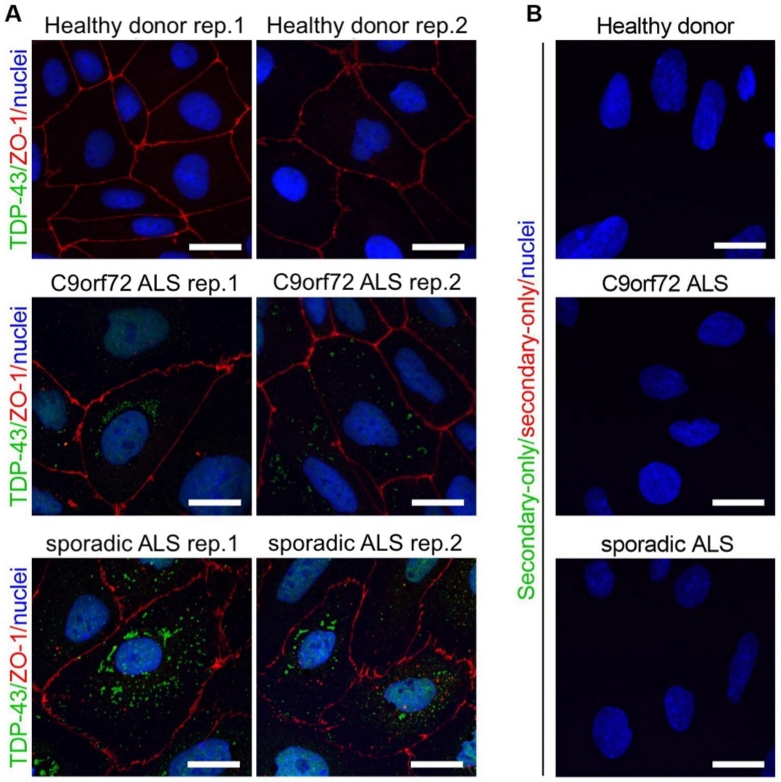

**Supplementary figure S3. Expression of TDP-43 protein in control and ALS iBECs and secondary-only controls for anti-TDP-43 and anti-ZO-1 antibodies.** (**A**) Additional examples of representative immunofluorescence images of TDP-43 (green) and ZO-1 (red) with Hoechst nuclear counterstaining (blue) in healthy donor, C9orf72 ALS, and sporadic ALS iBECs. (**B**) Representative immunofluorescence images of secondary antibody only controls for anti-TDP-43 (green) and ZO-1 (red) in healthy donor, C9orf72 ALS, and sporadic ALS iBECs. Nuclei (blue) stained with Hoechst. Scale bar = 20 µm.

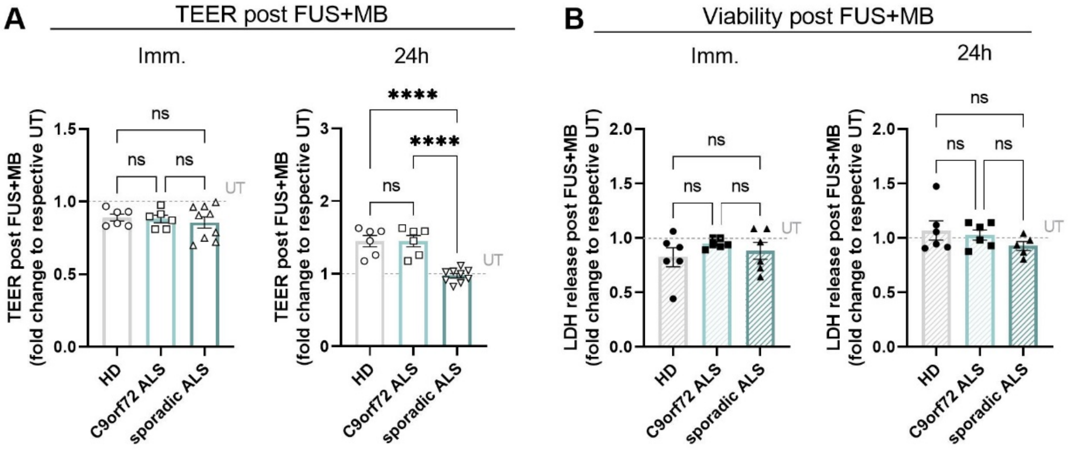

**Supplementary figure S4. Comparison of the effects of FUS$^{+MB}$ on control and ALS iBEC monolayer integrity and viability.** (**A**) Comparison of fold changes in TEER (Ohm x cm$^2$) of healthy donor, C9orf72 ALS and sporadic ALS iBEC exposed to FUS$^{+MB}$, immediately and 24 h following the treatment. TEER shown as fold changes to respective untreated (UT) cells at each time point. A minimum of $n$ = 6 independent replicates per line. (**B**) Comparison of fold changes in relative lactate dehydrogenase (LDH) release in healthy donor, C9orf72 ALS and sporadic ALS iBEC exposed to FUS$^{+MB}$, immediately and 24 h following the treatment. LDH release shown as fold changes to respective untreated (UT) cells at each time point. A minimum of $n$ = 5 independent replicates per line. Data analysed with One-way ANOVA with Tukey's test, error bars = SEM. ****$P$<0.0001.

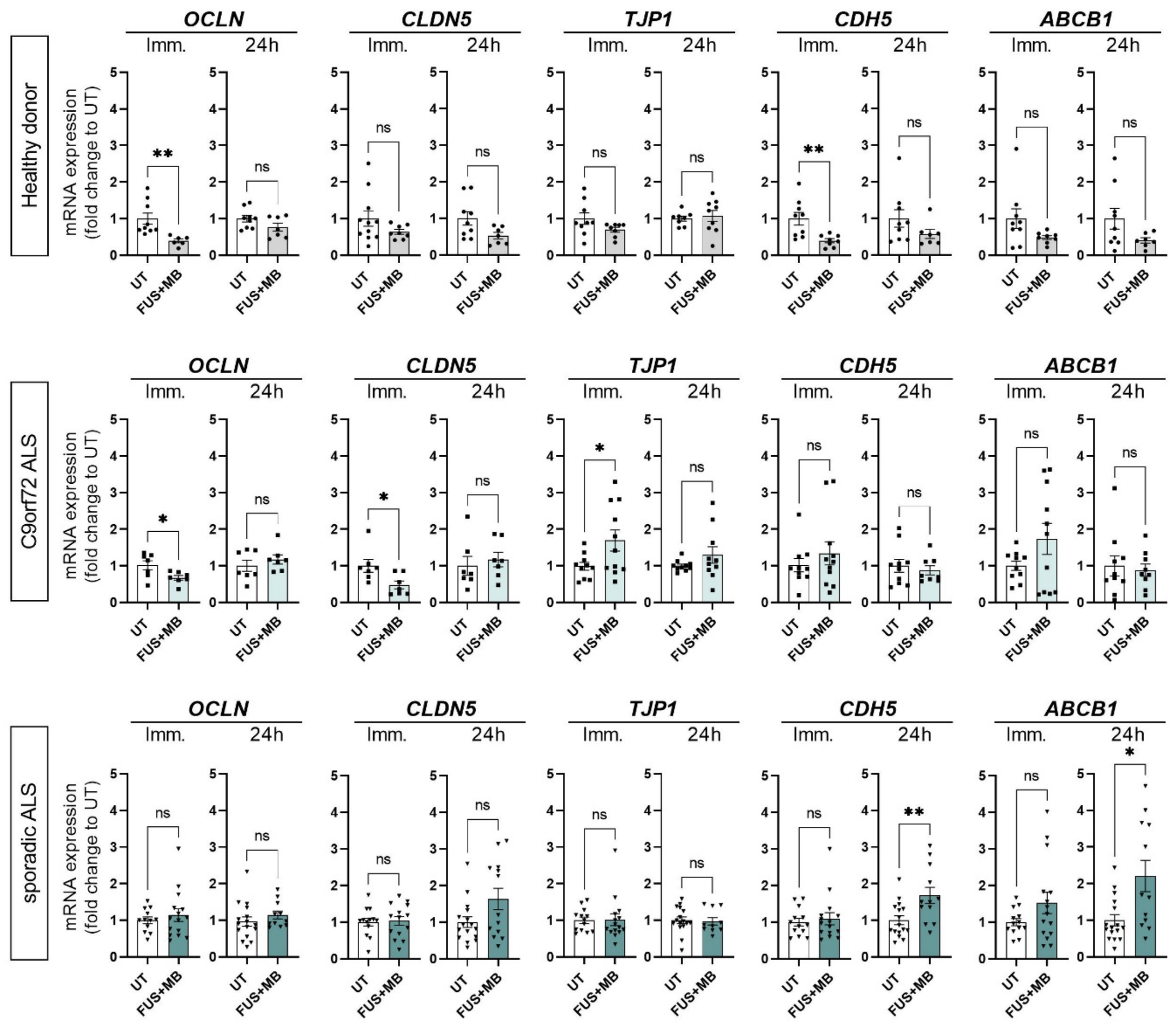

**Supplementary figure S5. Effects of FUS$^{+MB}$ on the expression of junctional and transporter marker genes in control and ALS iBECs.** Fold changes changes in mRNA expression of *OCLN*, *CLDN5, TJP1*, *CDH5*, *ABCB1* in healthy donor, C9orf72 ALS, and sporadic ALS iBEC exposed to FUS$^{+MB}$, immediately and 24 h following the treatment. Data presented as fold changes to respective untreated (UT) cells at each timepoint. A minimum of $n$ = 6 independent replicates per line. Data analysed with Student's $t$-test or Welch's $t$-test. *$P$<0.05, **$P$<0.01.

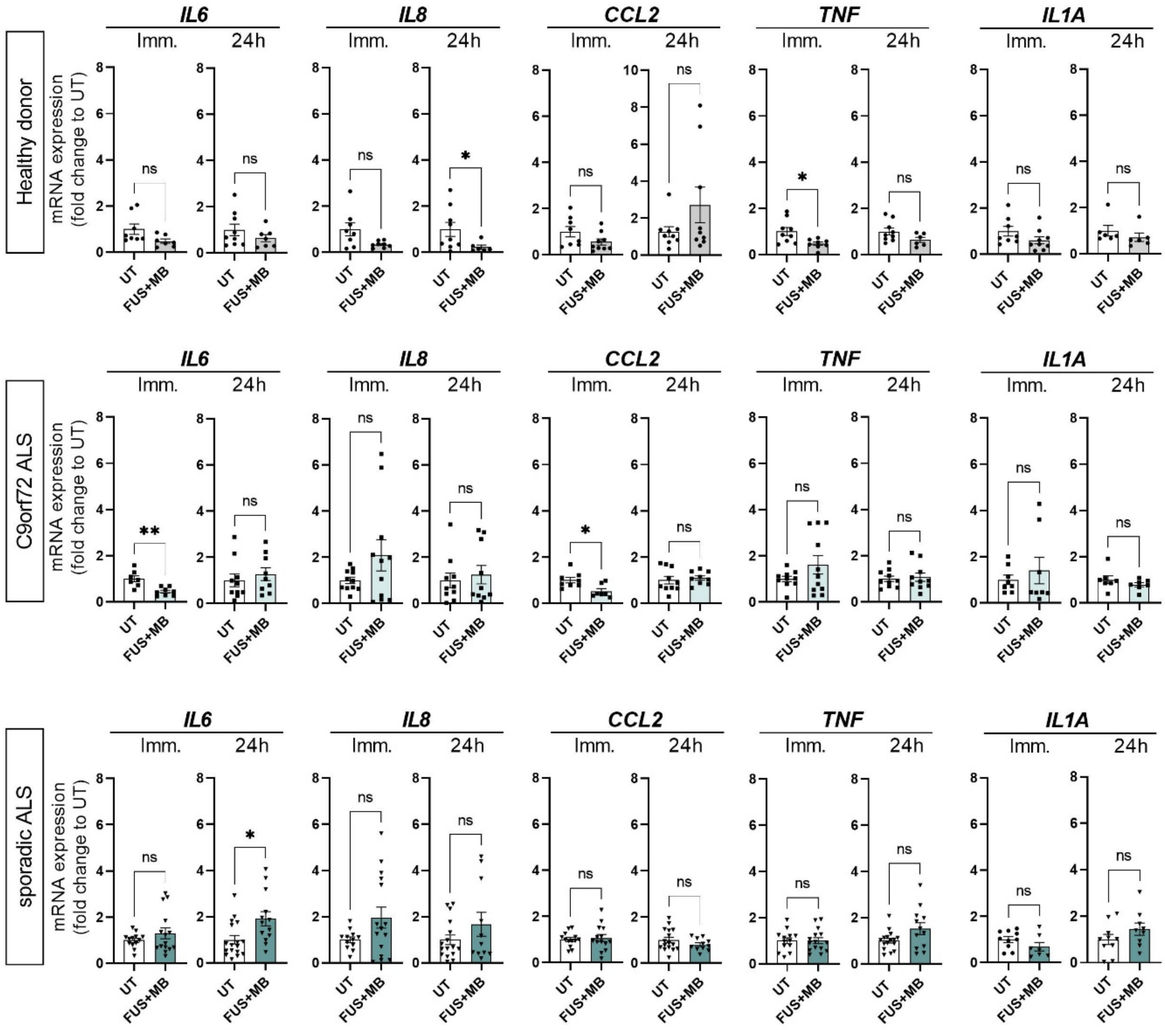

**Supplementary figure S6. Effects of FUS$^{+MB}$ on the expression of inflammatory marker genes in control and ALS iBECs.** Fold changes changes in mRNA expression of *IL6*, *IL8, CCL2*, *TNF* and *IL1A* in healthy donor, C9orf72 ALS, and sporadic ALS iBEC exposed to FUS$^{+MB}$, immediately and 24 h following the treatment. Data presented as fold changes to respective untreated (UT) cells at each timepoint. A minimum of $n$ = 6 independent replicates per line. Data analysed with Student's $t$-test or Welch's $t$-test. *$P$<0.05, **$P$<0.01.

**SUPPLEMENTAL TABLES:**

**Supplementary table S1**. Human iPSC lines used in the study.

| **Disease phenotype** | **Cell line ID** | **Age at collection** | **Gender** |
|---|---|---|---|
| Healthy donor (HD) control line | HDFa | Young adult (exact age unknown) | Female |
| C9orf72 carrier (familial ALS) | IT_004-04 | 30 | Male |
| Sporadic ALS | IT_002-04 | 39 | Female |

**Supplementary table S2**. Clinical and demographic characteristics of the ALS patients.

| **Parameter** | **Familial ALS patient (IT_004-04)** | **Sporadic ALS patient (IT_002-04)** |
|---|---|---|
| Age at onset (years) | 30 | 36 |
| Age at death/tracheostomy (years) | N.A. | 40 |
| Site of onset (spinal (S) or bulbar (B)) | N.A. | S |
| Survival (months) | N.A. | 49 |
| Gender | Male | Female |
| ΔFS[1] | N.A. | 0.78 |
| ALSFRS-R at entry[2] | 48 | 41 |
| Genetics | C9orf72 (> 30 GGGGCC) | None (sporadic) |

[1] **ΔFS score**: 48 − ALSFRS-R score at time of diagnosis) / interval (months) from onset to diagnosis. The score allows to identify three putative rates of progression: slow (ΔFS b 0.5), intermediate (ΔFS between 0.5 and 1.0), rapid (ΔFS > 1.0), which can predict survival [*F. Kimura, C. Fujimura, S. Ishida, et al., Progression rate of ALSFRS-R at time of diagnosis predicts survival time in ALS, Neurology 66 (2006) 265–267*]

[2] **ALSFRS-R**, range: 0 (complete paralysis, locked-in) – 48 (normal) [*J.M. Cedarbaum, N. Stambler, E. Malta, et al., The ALSFRS-R: a revised ALS functional rating scale that incorporates assessments of respiratory function. BDNF ALS study group (phase III), J. Neurol. Sci. 169 (1999) 13–21*]

**N.A.**: not applicable

**Supplementary table S3.** Antibodies used for the immunofluorescence analysis.

| **Primary antibodies** | **Species** | **Source** | **Identifier** | **Dilution** |
|---|---|---|---|---|
| SOX2 | rat | Invitrogen | Cat#14981182 | 1:100 |
| Nanog | rabbit | Abcam | Cat#ab21624 | 1:100 |
| ZO-1 | mouse | Invitrogen | Cat#339100 | 1:100 |
| occludin | rabbit | Invitrogen | Cat#711500 | 1:100 |
| claudin-5 | mouse | Invitrogen | Cat#352500 | 1:100 |
| Glut1 | mouse | Invitrogen | Cat#MA5-11315 | 1:100 |
| TDP-43 | rabbit | Proteintech | Cat#10782-2-AP | 1:200 |
| **Secondary antibodies** | **Species** | **Source** | **Identifier** | **Dilution** |
| anti-mouse Alexa Fluor 488 | goat | Invitrogen | Cat#A11029 | 1:250 |
| anti-mouse Alexa Fluor 594 | goat | Invitrogen | Cat#A11032 | |
| anti-mouse Alexa Fluor 647 | goat | Invitrogen | Cat#A32728 | |
| anti-rabbit Alexa Fluor 488 | goat | Invitrogen | Cat#A11034 | |
| anti-rat Alexa Fluor 647 | goat | Invitrogen | Cat#A21247 | |

**Supplementary table S4.** Primer sequences used in the study.

| **Target gene** | **Forward primer sequence** | **Reverse primer sequence** |
|---|---|---|
| (Sex determining region Y)-box 2 (*SOX2*) | CCACCTACAGCATGTCCTACTCG | GGGAGGAAGAGGTAACACAGG |
| Nanog homeobox *(NANOG)* | ACCTCAGCTACAAACAGGTGAA | AAAGGCTGGGGTAGGTAGGT |
| Octamer-binding transcription factor 4 (*OCT4)* | ATCTTCAGGAGATATGCAAAGCAGA | TGATCTGCTGCAGTGTGGGT |
| VE-cadherin *(CDH5)* | AGGCAAGATCAAGTCAAGCGT | GAGTCTCCAGGTTTTCGCCA |
| Claudin-5 *(CLDN5)* | GATTGAGAGGTCTGGGAAGCC | ATCCCATGGCAAACAGAGAGG |
| Occludin *(OCLN)* | GAAGCAAGTGAAGGGATCTGC | ACAACTTGGCATCAGCCTTCT |
| Zonula occludens-1 *(TJP1)* | ACAGCTACAGGAAAATGACCGA | ACTGGTTCAGGATCAGGACG |
| P-glycoprotein (*ABCB1*) | CAGATAAAAGAGAGGTGCAACGG | GCCCGGATTGACTGAATGCT |
| Interleukin-6 *(IL6)* | TGCAATAACCACCCCTG ACC | TGCGCAGAATGAGATGAGTTG |
| Interleukin-8 *(IL8)* | AGACAGCAGAGCACACA AGC | ATGGTTCCTTCCGGTGGT |
| Interleukin-1A (*IL1A*) | CATCGCCAATGACTCAGAGAAG | TGCCAAGCACACCCAGTAGTCTTGCTT |
| Monocyte chemoattractant protein-1 *(CCL2)* | GCTCATAGCAGCCACCTTCATTC | GGACACTTGCTGCTGGTGATTC |
| Tumour necrosis factor α (*TNF*) | CAGCCTCTTCTCCTTCC TGAT | GCCAGAGGGCTGATTAGAGA |
| *18S* | TTCGAGGCCCTGTAATTGGA | GCAGCAACTTTAATATACGCTATTGG |
